# NHSL3 interacts with Ena/VASP proteins and the Scar/WAVE complex and promotes cell migration

**DOI:** 10.1101/2025.04.03.647056

**Authors:** Tommy Pallett, Fuad Mosis, Simon Poland, Simon M. Ameer-Beg, Matthias Krause

**Affiliations:** King’s College London, Krause group, Randall Centre for Cell and Molecular Biophysics, New Hunt’s House, Guy’s Campus, London, SE1 1UL, UK; King’s College London, Ameer-Beg group, Richard Dimbleby Cancer Research Laboratories, Comprehensive Cancer Centre, School of Cancer and Pharmaceutical Sciences, New Hunt’s House, Guy’s Campus, London, SE1 1UL, UK; King’s College London, Poland group, Comprehensive Cancer Centre, School of Cancer and Pharmaceutical Sciences, New Hunt’s House, Guy’s Campus, London, SE1 1UL, UK

## Abstract

Tight control of mesenchymal cell migration is important for embryonic development and its deregulation causes disease. It is driven by lamellipodia protrusion, the leading edge of the migrating cell. This is controlled by Rac-Scar/WAVE-Arp2/3 complexes driving actin filament nucleation coupled to Ena/VASP proteins mediating actin filament elongation. These activities are coordinated by leading-edge proteins, such as Lamellipodin and NHSL1. Here, we discovered KIAA1522/NHSL3 as an additional regulator of these essential actin effectors. We reveal that NHSL3 promotes cell migration. NHSL3 co-localises at the edge of lamellipodia with Ena/VASP proteins and the Scar/WAVE complex. We show that it binds to Ena/VASP proteins and the Scar/WAVE complex and functions to inhibit Scar/WAVE-Arp2/3 activity in cells. NHSL3 interacts with the Scar/WAVE complex subunit Abi and, in contrast to other known Scar/WAVE complex binders, additionally to the CYFIP1/2 subunit through 3 short linear motifs. Thus, control of actin filament nucleation and elongation at the leading edge of mesenchymal cells is more complex than anticipated. Our study provides insights into the intricate regulation of lamellipodial actin networks highly relevant for understanding control of mesenchymal cell migration during development and diseases.

## Introduction

Cell migration is essential during embryonic development as well as for adult homeostasis in processes such as immune response, wound healing, and tissue repair. Its deregulation can therefore lead to developmental defects or delayed wound healing for example^1–3^, and the dysregulation of migration can drive metastasis during cancer progression^4^. There are various modes of migration, both single-cell and collective. For example, fibroblasts, which are required for wound healing, migrate as single cells in a mesenchymal mode. Similarly, cancer cells, which are characteristically plastic, can also migrate using this mode after undergoing an epithelial to mesenchymal transition^5,6^. Mesenchymal migration is characterised by an actin dependent protrusion, the lamellipodium, at the leading edge of the cell, which is stabilised by integrin dependent adhesions to the extracellular matrix^7–9^. Branched F-actin nucleation induced by the Arp2/3 complex and subsequent polymerisation of the filaments directly underneath the plasma membrane provides the force for lamellipodium extension^10–12^.

The Scar/WAVE complex, also known as the WAVE Regulatory Complex (WRC), is heteropentameric, composed of CYFIP1-2, Nap1, Scar/WAVE1-3, Abi1-3, and HSPC300. Scar/WAVE proteins harbour a C-terminal WCA region which binds and activates the Arp2/3 complex^13–17^. This interface is inaccessible in the autoinhibited conformation of the Scar/WAVE complex and this autoinhibition is relieved by active-Rac, tyrosine phosphorylation, and phosphoinositide binding^15,18–20^. The same signals also localise Lamellipodin to the edge of lamellipodia which binds via three proline-rich motifs to the SH3 domain of the Abi subunit in the Scar/WAVE complex. Both, Lamellipodin and the Scar/WAVE complex are required for lamellipodium formation and Lamellipodin promotes mesenchymal cell migration via its interaction with the Scar/WAVE complex^21–25^.

Lamellipodin also recruits critical F-actin elongation factors of the Ena/VASP family of proteins^21,22,25^. The Ena/VASP protein family comprises VASP, Mena, and EVL in vertebrates^26,27^. All family members contain an N-terminal EVH1 domain which can bind to proline-rich regions of other proteins. EVH1 domain binding motifs consist of a core proline-rich sequence F/Y/W/LPXΦP (where X is any and Φ is a hydrophobic amino acid) flanked by acidic amino acids^28–31^. Lamellipodin interacts with Ena/VASP proteins through seven such motifs^21^ Ena/VASP proteins function to increase F-actin filament elongation by recruiting polymerisation-competent Profilin-bound monomeric actin and temporarily protecting the F-actin filament ends from capping proteins^32–38^. Lamellipodin therefore may act as a coordinator of Scar/WAVE-Arp2/3 complex-mediated lamellipodial protrusion and Ena/VASP-mediated F-actin elongation^7^.

The regulation of F-actin filament branching and elongation is more complex than previously anticipated. We identified Nance-Horan Syndrome-like 1 (NHSL1) protein as another interactor of the Scar/WAVE complex at the leading edge of cells, but one which negatively regulates Scar/WAVE-Arp2/3 complex activity and thus functions as an inhibitor of mesenchymal cell migration^39^. NHSL1 also binds directly to the Scar/WAVE complex via the SH3 domain of the Abi subunit using, in this case, two SH3 domain binding motifs^39^. Recently, we found that NHSL1 additionally interacts with Ena/VASP proteins^40^ implying that, like Lamellipodin, it is also able to coordinate both branching and elongation within the lamellipodium.

NHSL1 is part of a protein family comprised of Nance-Horan Syndrome protein (NHS), NHSL1, and NHSL2^41^. Nance-Horan syndrome is a rare X-linked developmental disorder characterised by cataracts, facial dysmorphism, dental abnormalities, and cognitive impairment caused by mutations in the *nhs* gene^42,43^.

Obtaining tightly controlled protrusion in mesenchymal cell migration must require a high level of coordination between the activities of the Scar/WAVE-Arp2/3 complexes and Ena/VASP proteins to balance the branching and elongation of F-actin. We therefore hypothesised that to fully understand this biological process, additional regulators of these essential actin effectors needed to be identified. In this study, we identified the previously poorly characterised protein KIAA1522^44^ as another regulator of both the Scar/WAVE complex and Ena/VASP. Human *KIAA1522* and its mouse ortholog C77080 are highly conserved across vertebrate species (Ensembl ID: ENSG00000162522). Overexpression of KIAA1522 has been linked to several cancers including lung, oesophageal, and liver^45–47^. We considered that KIAA1522 could be the fourth member of the NHS protein family, due to similarities in biochemistry and functional roles in the cell. Since then, KIAA1522 has been formerly recognised by HUGO as NHSL3 (HGNC ID: 29301) and we henceforth refer to it as such.

We created a NHSL3 CRISPR knockout cell line which revealed that NHSL3 promotes cell migration efficiency. As with NHSL1, NHSL3 localises the very edge of lamellipodia where it co-localises with Ena/VASP proteins and the Scar/WAVE complex. We show that it binds to the Ena/VASP proteins, Mena and VASP, through an extended EVH1 domain binding motif. We found that NHSL3 also interacts with the Scar/WAVE complex by directly binding to the Abi subunit, but in this case the interaction is mediated by only one SH3 domain binding site. The NHSL3-Scar/WAVE complex interaction functions similarly to NHSL1 by inhibiting Scar/WAVE-Arp2/3 activity - a surprising result, given that NHSL3 promotes migration efficiency. Unexpectedly, and in contrast to Lamellipodin and NHSL1, we discovered that NHSL3 displays another extended interface, comprised of 3 short linear motifs, which mediates an interaction with the CYFIP1/CYFIP2 subunits of the Scar/WAVE complex. Overall, these results suggest that the control of protrusion in mesenchymal cells through the interplay of actin filament nucleation and elongation is incredibly complex, and that simple linear relationships between variables such as Arp2/3 activity and migration efficiency do not represent the reality of the processes involved. Our study provides insights into this intricate regulation which is highly relevant and necessary for understanding control of mesenchymal cell migration during development and disease.

## Results

We identified the poorly characterised protein KIAA1522 as a putative interactor of the EVH1 domain of Ena/VASP proteins by performing a BLAST search of GenBank for EVH1 domain binding motifs. One of the hits, KIAA1522, harbours one canonical EVH1 domain binding motif F/Y/W/LPXΦP flanked by acidic amino acids followed by a second putative non-canonical site which lacks the fourth proline residue^28,29^ (705-D-p-s-W-P-P-P-P-P-P-a-P-E-E-q-D-l-s-m-a-D-F-P-P-P-E-E-731; *Mus musculus* KIAA1522). KIAA1522 is widely expressed in diverse mouse tissues as a transcript of around 5.5kb (Fig. 1A) which agrees with expression detected in human tissues from RNAseq data available from the Human Protein atlas (NCBI Gene ID: 57648, Human; 97130, *Mus musculus*). Only exon 1 out of the seven exons is alternatively spliced (1a-c) resulting in the expression of at least 3 different isoforms (KIAA1522 isoform 1a-1c) (Fig. 1D).

**Figure 1.**
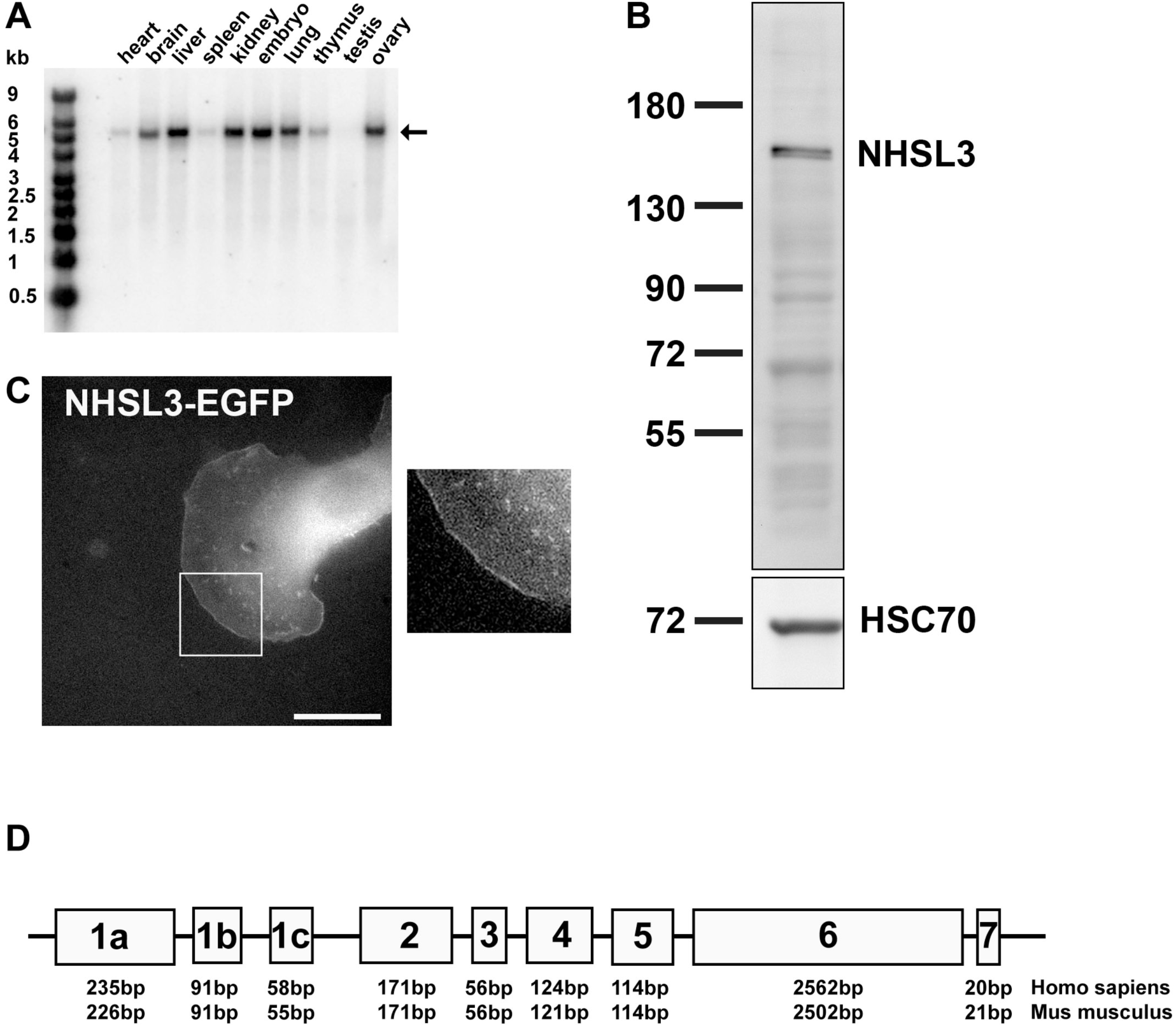
NHSL3 localises to the edge of lamellipodia and to vesicular structures. **(A)** Northern blot analysis of NHSL1 expression in different tissues. The arrow shows a large 5.5kb cDNA that is ubiquitously expressed but low in heart and spleen and not in testis. **(B**) Western blots detecting NHSL3 protein in the B16-F1 melanoma cell line using the polyclonal antibody (3915); HSC70 serves as the loading control. **(C)** C-terminal EGFP-tagged NHSL3 was expressed in B16-F1 cells and, after plating on laminin, localisation was imaged live. Representative images shown from three independent experiments. Scale bar: 20 μm. Inset represents a magnified view of the white box. See also related video S1. **(D)** Intron and exon structure of NHSL3 isoforms is depicted with exon size in base pair length (bp) for Homo sapiens or Mus musculus below.

**Figure S1.**
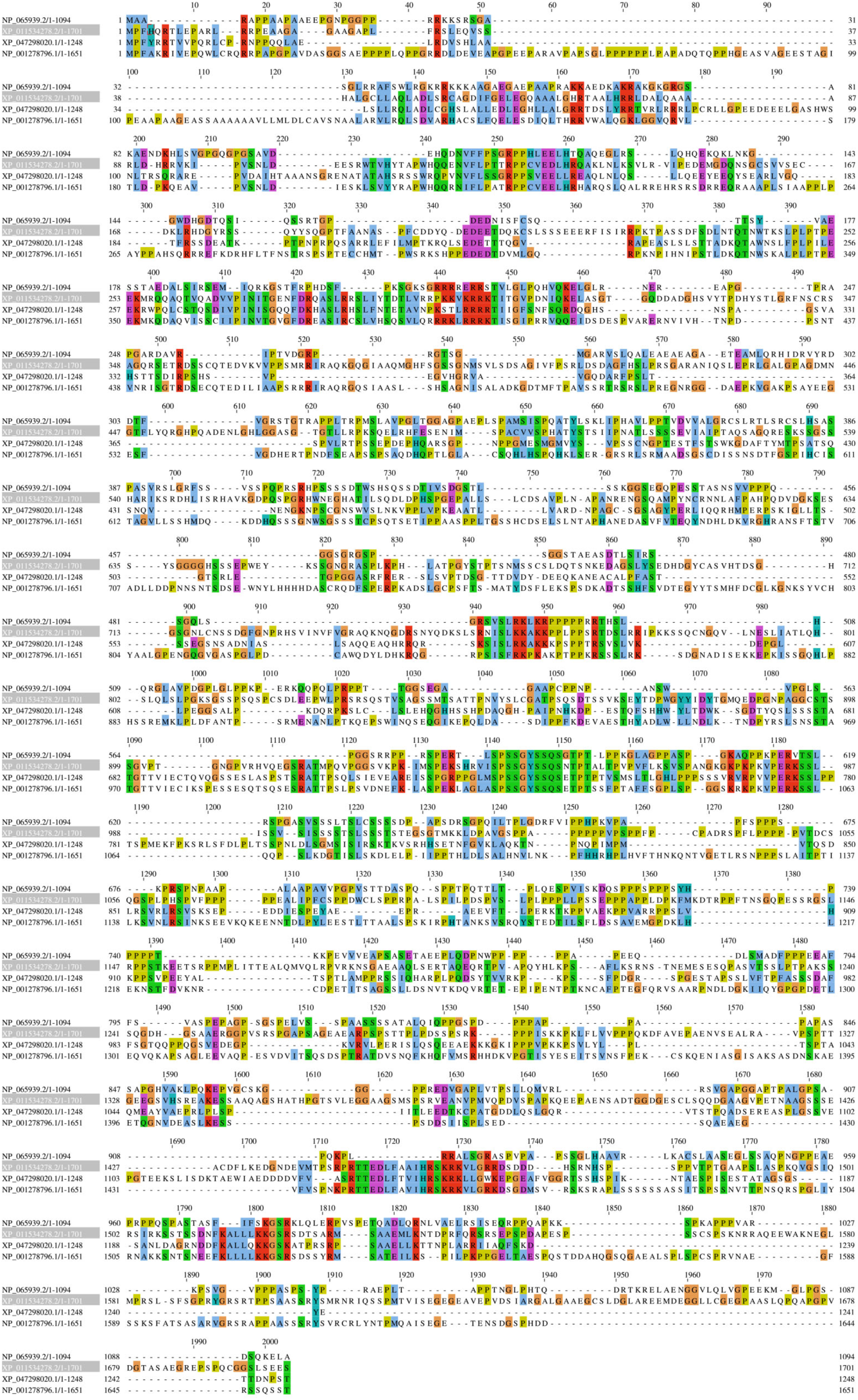
KIAA1522/NHSL3 is a member of the NHS family. **(A)** Multiple Alignment using Fast Fourier Transform (MAFFT) was performed on the EBI server (https://www.ebi.ac.uk/jdispatcher/msa/mafft?stype=protein) using proteins sequences of human NHS (NP_001278796.1), NHSL1 (XP_011534278.2), NHSL2 (XP_047298020.1), and NHSL3/KIAA5122 (NP_065939.2).

We cloned the full-length mouse KIAA1522 isoform 1c from EST clone BE309569 adding the missing first 10 amino acids in exon 1c (1013 aa total). As previously mentioned, KIAA1522 has recently been renamed by HUGO as NHSL3, and we will now refer to it as such. NHSL3 in human is only 29.3% conserved at the amino acid level compared to NHSL1, however this is not dissimilar to the conservation between other members of the NHS family (Fig S1). The mouse NHSL3 isoform c as a cDNA is called NHSL3 Transcript Variant 3 (NM_001285865.2) and is equivalent to the human NHSL3 Transcript Variant 2 (NM_001198972.2). All future references of NHSL3 in the context of the assays carried out in this study refer to mouse NHSL3 isoform 1c unless otherwise specified.

### NHSL3 co-localises with Mena and the Scar/WAVE complex at the very edge of lamellipodia

To explore the localisation and function of NHSL3, we generated a rabbit polyclonal antiserum against a fragment of NHSL3 (aa535-701; GST-4 see Fig. 5E). This antiserum detected a protein of 170 kDa in lysates of the B16-F1 melanoma cell line (Fig. 1B). NHSL3 C-terminally tagged with EGFP (NHSL3-EGFP) localised to the very edge of lamellipodia and to vesicular structures emanating from it towards the cytosol (Fig. 1C; Supplemental movie 1). This is reminiscent of the localisation of NHSL1 and NHSL2 that we observed previously^39,40^.

To explore potential co-localisation with Ena/VASP proteins, which are known to localise to the leading edge of migrating cells, to the tips of microspikes, and to focal adhesions, we co-expressed NHSL3-EGFP with mScarlet-Mena in B16-F1 cells and observed that both proteins co-localise at the very edge of lamellipodia but not at microspikes or focal adhesions (Fig. 2A; Supplemental movie 2). Since the Scar/WAVE complex also resides at the very edge of lamellipodia and interacts with the other NHS family members NHS and NHSL1, we co-expressed NHSL3-EGFP with mScarlet tagged Scar/WAVE1, 2, or 3 or the Scar/WAVE complex components Abi1 or Nap1. We observed co-localisation of all the Scar/WAVE complex components with NHSL3 at the very edge of lamellipodia (Fig. 2B-D; Fig. S2A,B; Supplemental movies 3-7). To verify that this co-localisation is not due to tagging or overexpression, we performed immunofluorescence and observed a similar localisation of endogenous NHSL3 to the very edge of lamellipodia and to vesicular structures. Again, we found that endogenous NHSL3 co-localised with Mena, Scar/WAVE1, and Abi1 at the very edge of lamellipodia (Fig. 3A-C).

**Figure 2.**
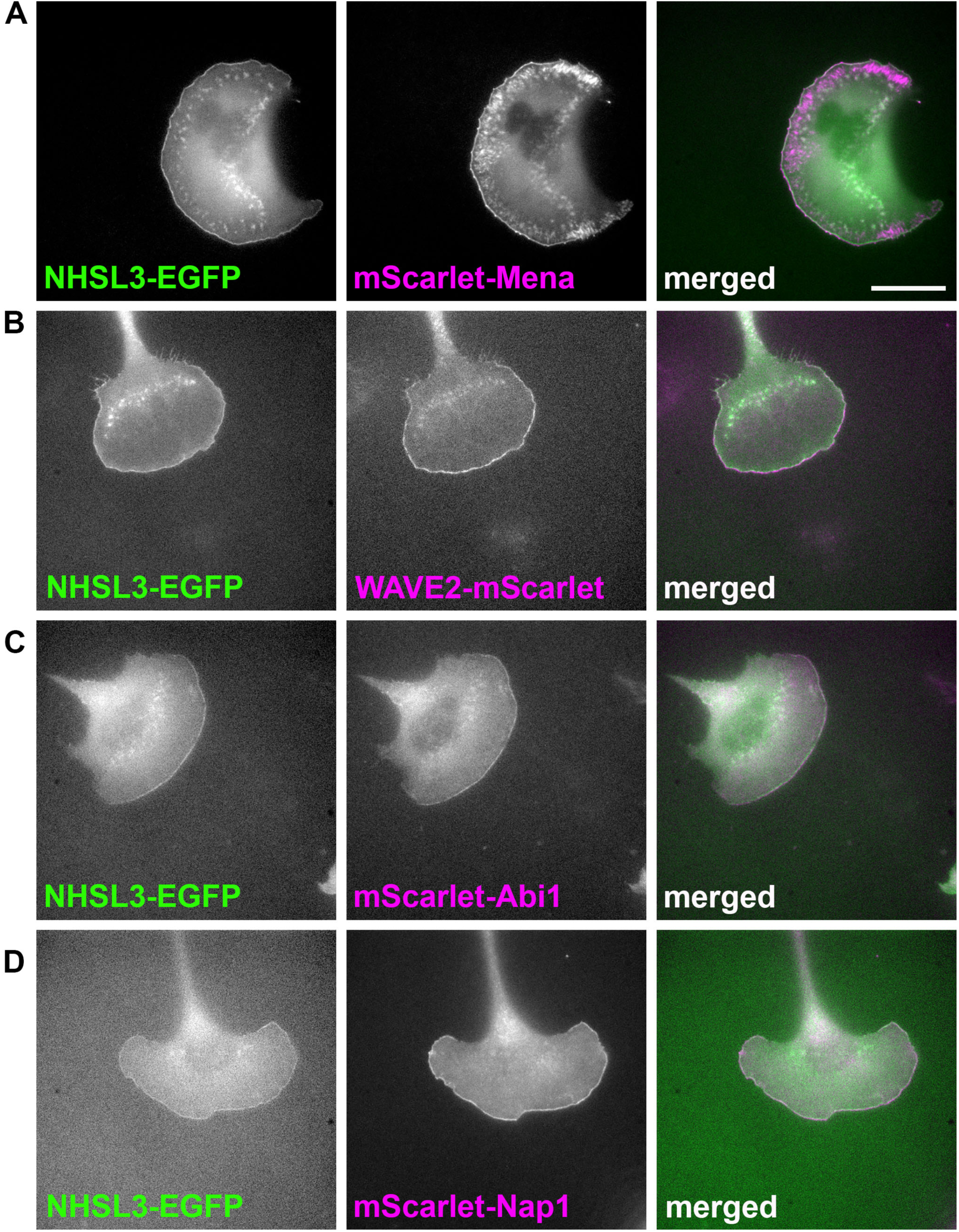
NHSL3 co-localises with Mena and the Scar/WAVE complex at the edge of lamellipodia. **(A-D)** Still images from live cell imaging showing NHSL3-EGFP co-expressed with mScarlet-I-tagged Mena **(A)**, -Scar/WAVE2 **(B)**, -Abi1 **(C)** and **-**Nap1 **(D)** in B16-F1 cells. Representative images shown from three independent experiments. Scale bar in (A) for (A-D) represent 20 µm. See also related videos S2-5.

**Figure S2.**
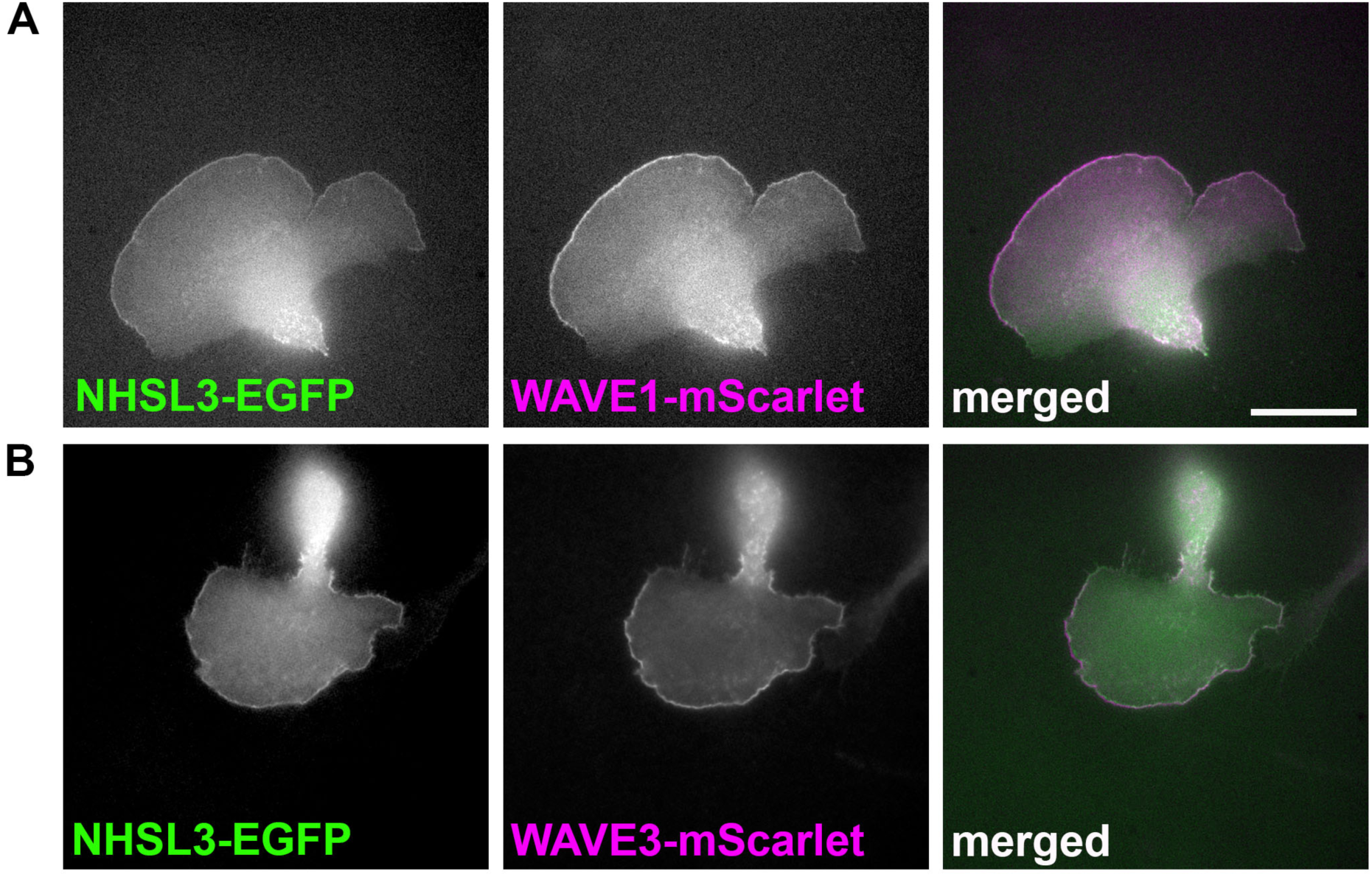
NHSL3 co-localises with the Scar/WAVE complex at the edge of lamellipodia. **(A,B)** Still images from live cell imaging showing NHSL3-EGFP co-expressed with mScarlet-I-tagged -Scar/WAVE1 **(A)**, -Scar/WAVE3 **(B)** in B16-F1 cells. Representative images shown from three independent experiments. Scale bar in (A) for (A,B) represent 20 µm. Data relates to Fig. 2. See also related videos S6-7.

**Figure 3.**
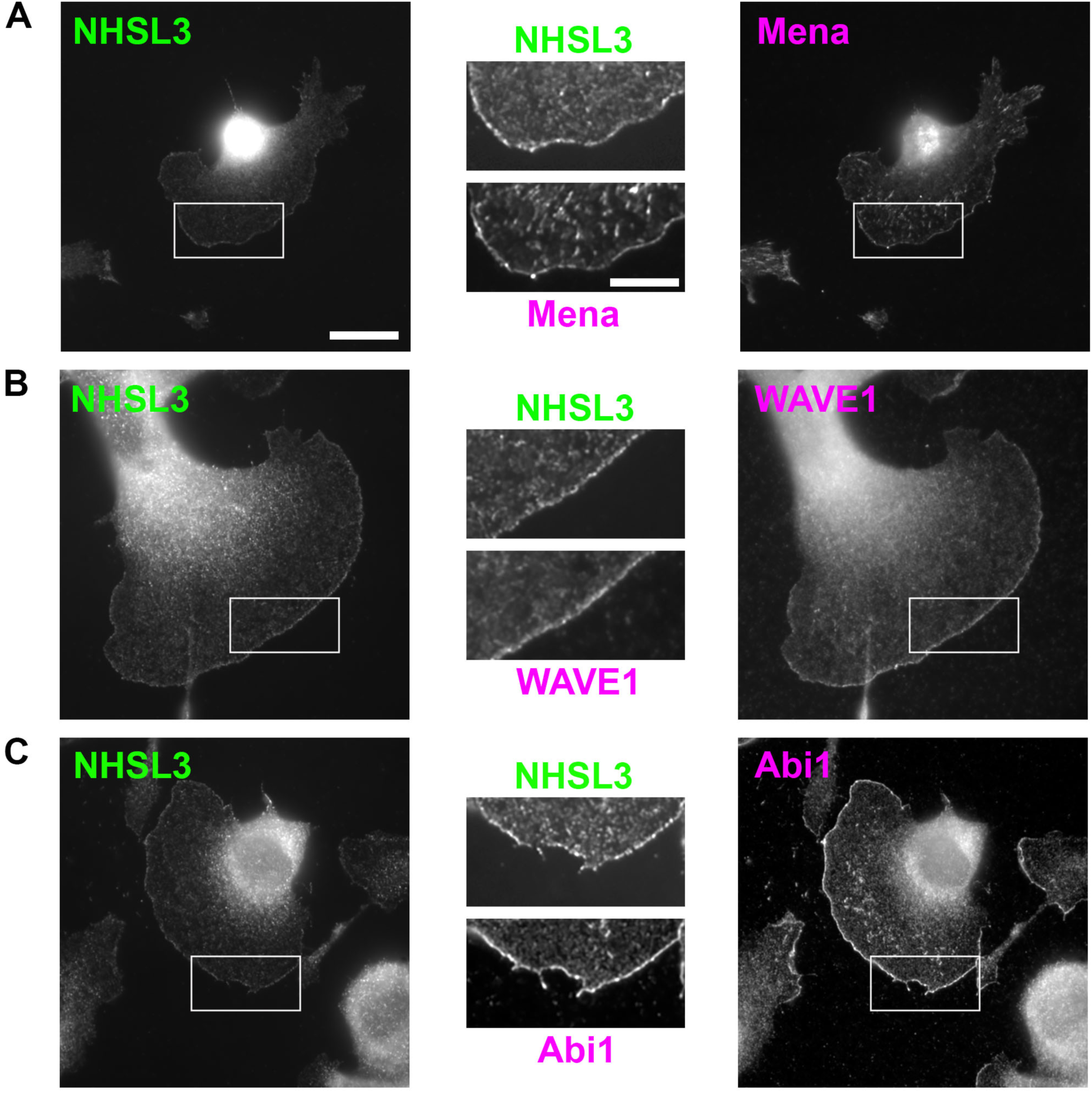
Endogenous NHSL3 co-localises with Mena and the Scar/WAVE complex at the edge of lamellipodia. **(A-C)** Endogenous NHSL3 co-localises with Mena (A, NHSL3 pAb; Mena mAb), Scar/WAVE1 (B, NHSL1 pAb; Scar/WAVE1 mAb), and Abi1 (C, NHSL1 pAb; Abi1 mAb) at the edge of lamellipodia in B16-F1 mouse melanoma cells. Scale bar in (A) applies also to (B,C): 20 µm. Inset represents a magnified view of the white box. Scale bar in inset in (A) applies also to inset for (B,C): 20 μm. Representative images shown from three independent experiments.

### NHSL3 is a novel interactor of Ena/VASP proteins

To investigate the potential interaction with Ena/VASP proteins, we co-transfected HEK cells with NHSL3-EGFP and Myc-tagged VASP, Mena, or EVL. We pulled-down NHSL3-EGFP from cell lysates and found that NHSL3 interacts with all three Ena/VASP proteins (Fig. 4A,B). To explore whether the interaction is mediated by the EVH1 domain in Ena/VASP proteins, we purified this domain tagged with GST from *E. coli* (Fig. 4E). Bead immobilised, purified GST-EVH1 domains from VASP, Mena, or EVL were used to pulldown NHSL3-EGFP from lysates of transfected HEK cells. This revealed that NHSL3 most strongly bound to the EVH1 domain of Mena, more weakly to VASP and not detectably to EVL (Fig. 4D). This result was at first surprising considering we observed interaction with all three Ena/VASP proteins in co-pulldowns (Fig. 4A,B). However, Ena/VASP proteins are known to hetero-oligomerise ^48^, and so the interactions of NHSL3 with specific Ena/VASP proteins may be co-dependent. Mena and EVL show only weak hetero-oligomerisation ^48^, suggesting that NHSL3 may specifically mediate part of its function through Mena. However, since VASP heterotetramerises without bias, the co-pulldown with EVL (Fig. 4A,B) may be the result of an indirect interaction. We next performed endogenous co-immunoprecipitation experiments using our antibody against NHSL3, observing that endogenous NHSL3 associates with Mena (Fig. 4C).

**Figure 4.**
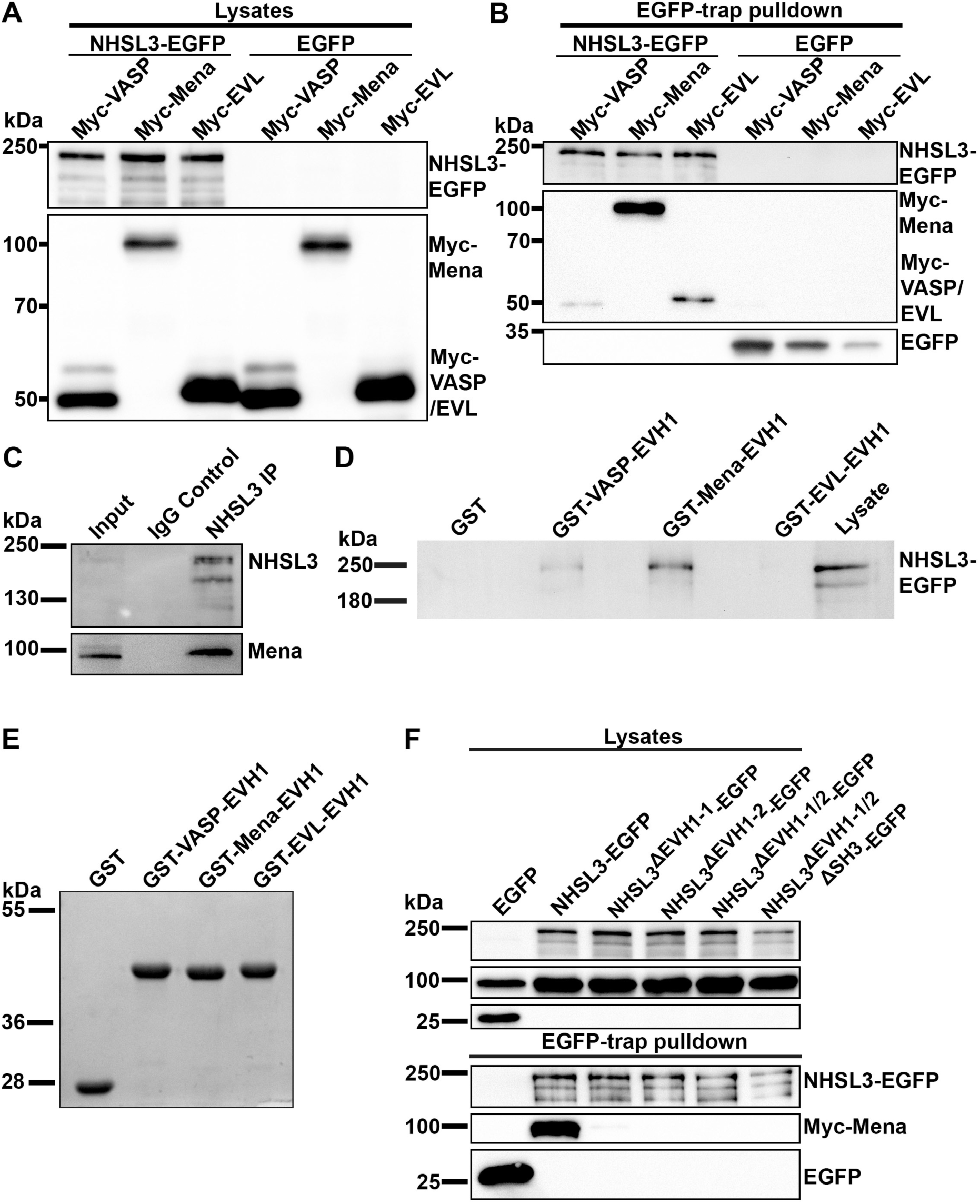
NHSL3 interacts with Ena/VASP proteins via two FP4 motifs. (**A-B**) Myc-VASP, Myc-Mena or Myc-EVL were co-expressed with NHSL3-EGFP or EGFP alone in HEK cells. EGFP-tagged NHSL3 was pulled down from cell lysates (A) using a nanobody against EGFP (B) and blots were probed using antibodies against Myc and EGFP. Blot representative of four independent experiments. (**C**) Endogenous NHSL3 was immunoprecipitated with NHSL3 polyclonal antiserum or normal rabbit IgG as negative control from B16-F1 lysates, blotted and probed using antibodies against Mena and NHSL3. Blot representative of 3 independent experiments. (**D**) NHSL3-EGFP was expressed in HEK cells. Purified GST-tagged EVH1 domains, or GST only (E: Coomassie gel) as negative control, were used to pull down associated proteins from HEK lysates and blots were probed using an antibody against EGFP. Blots representative of three individual experiments. **(E)** Coomassie stained SDS-PAGE gels depicting purified GST or GST-tagged EVH1 domains from VASP, Mena, or EVL. (**F**) Myc-tagged Mena was co-expressed with EGFP only as control or EGFP-tagged wild-type NHSL3, EGFP-tagged NHSL3 with the first or second FP4 motif mutated (FP4-1 Mutant or FP4-2 Mutant), with both FP4 motifs mutated (FP4-1/2 Mutant), or with both FP4 motifs and the Abi SH3 domain binding site mutated (FP4-1/2+SH3 Mutant). Transfected EGFP-tagged proteins were pulled down from lysates using a nanobody against EGFP and blots were probed using antibodies against Myc and EGFP. Blot representative of four independent experiments.

**Figure S3.**
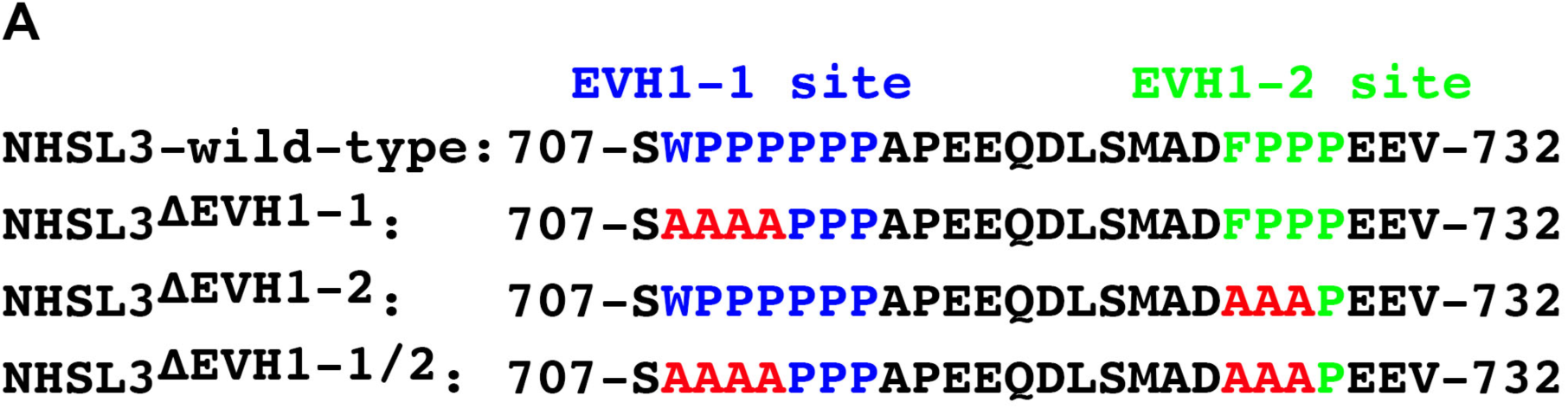
NHSL3 is a binding partner of Ena/VASP proteins. **(A)** The sequence of the two consecutive Ena/VASP EVH1 domain binding sites in NHSL3: EVH1 site 1 (EVH1-1 shown in blue) or EVH1 site 2 (EVH1-2 shown in green) and respective mutations (shown in red) introduced in the EVH1 site 1 (NHSL3**^ΔEVH1–^**), in the EVH1 site 2 (NHSL3**^ΔEVH1–2^**), or both sites (NHSL3**^ΔEVH1–/1^**) are shown. The amino acid numbering refers to murine NHSL3 isoform c (mmNHSL3 transcript variant 3 with exon1c; Genbank No: NM_001285866). Data relates to Fig. 4.

To explore whether the interaction is mediated through the putative EVH1 domain binding sites in NHSL3, we mutated these motifs either individually or together (Fig. S3A). We co-transfected these individual or double NHSL3-EGFP mutants (NHSL3ΔEVH1 -1, -2, or -1/2-EGFP) and wild-type NHSL3-EGFP as control together with Myc-Mena into HEK cells. EGFP pulldowns from cell lysates revealed that all three mutants were completely impaired in their interaction with Mena (Fig. 4F). This indicates that the interaction is mediated by the EVH1 domain of Mena and VASP and both EVH1 domain binding motifs in NHSL3 are required for this interaction.

### NHSL3 binds to the Scar/WAVE complex via the Abi SH3 domain

Since NHSL3 co-localises with the Scar/WAVE complex at the edge of lamellipodia, we investigated whether they interact with each other. We co-transfected HEK cells with NHSL3-EGFP and all Myc-tagged members of the Scar/WAVE complex: CYFIP1, Nap1, Scar/WAVE2, Abi1, and HSPC300. NHSL3 immunoprecipitation showed that NHSL3 interacts with all members of the Scar/WAVE complex when co-expressed (Fig. 5A). To verify that the endogenous proteins also form a complex, we immunoprecipitated endogenous NHSL3 and found that it indeed interacts with the Scar/WAVE complex components Scar/WAVE2 and Abi1 (Fig. 5B,C).

**Figure 5.**
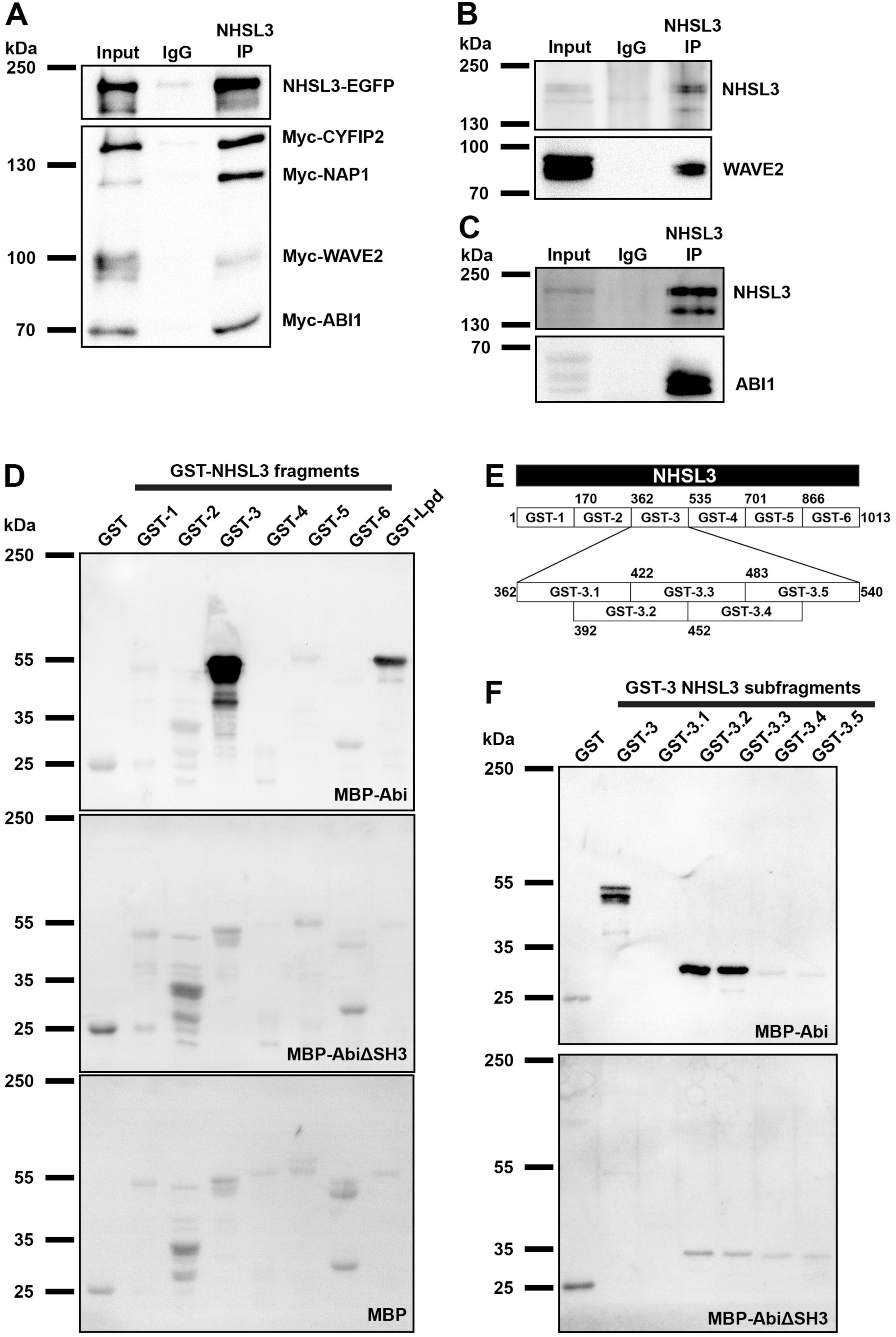
NHSL3 interacts with the Scar/WAVE complex via the Abi SH3 domain. (**A**) Myc-tagged CYFIP2, Nap1, Abi1, Scar/WAVE2, and HSPC300 were co-expressed with NHSL3-EGFP in HEK cells. EGFP-tagged NHSL3 was immunoprecipitated from cell lysates using NHSL3 polyclonal antiserum or normal rabbit IgG as negative control and blots were probed using antibodies against Myc and EGFP. Blot representative of three independent experiments. (**B,C**) Endogenous NHSL3 was immunoprecipitated with NHSL3 polyclonal antiserum or normal rabbit IgG as negative control from B16-F1 lysates, blotted and probed using antibodies against Scar/WAVE2 (B) or Abi1 (C) and NHSL3. Blot representative of 3 independent experiments. (**D-F**) Far western blot experiments using (D) six GST-tagged NHSL3 fragments (Fragment 1 - 6 in (E)) or (F) five sub-fragments of GST-3 (sub-fragment GST-3.1-3.5 in (E)) overlaid with purified MBP-Abi1 or MBP-Abi1ΔSH3 or MBP as negative control. GST-fusion protein containing several Abi SH3 binding motifs of the Lamellipodin protein^21^ served as a positive control and GST alone as the negative control. For (D,F) four independent experiments. **(E)** Schematic overview of the six GST-tagged fragments covering the entire amino acid sequence of NHSL3 and overlapping sub-fragments of fragment 3. The numbering refers to the amino acid numbers at the start and end of each fragment referring to murine NHSL3 isoform c (mmNHSL3 transcript variant 3 with exon1c; Genbank No: NM_001285866).

**Figure S4.**
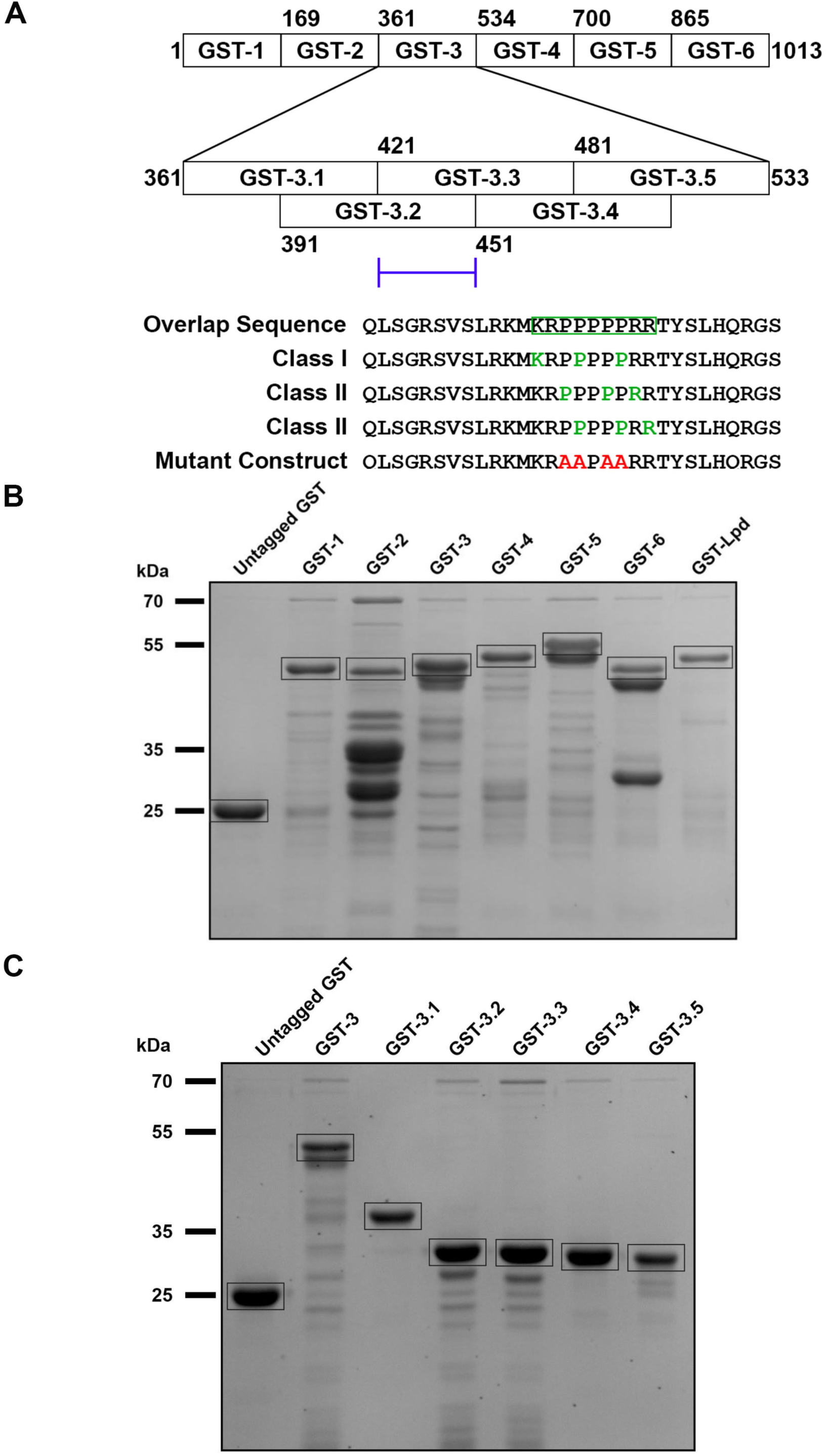
Coomassie gels of purified GST-tagged NHSL3 fragments. **(A)** Schematic overview of the six GST-tagged fragments covering the entire amino acid sequence of NHSL3 and overlapping sub-fragments of fragment 3. The numbering refers to the amino acid numbers at the start and end of each fragment referring to murine NHSL3 isoform c (mmNHSL3 transcript variant 3 with exon1c; Genbank No: NM_001285866). The location of the Abi SH3 domain binding site in GST-3.2 and GST-3.3 is indicated with a blue bracket. Below: Amino acid sequence of the overlap sequence harbouring the Abi SH3 domain binding site is shown with the amino acids in green representing class I or class II SH3 domain binding sites. The mutated amino acids are shown in red. **(B,C)** Coomassie gel of purified (B) GST and the six GST-tagged NHSL3 fragments and a GST-fusion protein containing several Abi SH3 binding motifs of the Lamellipodin protein^21^ as a positive control (Fragment 1 - 6 in (A)) or (C) five sub-fragments of GST-3 (sub-fragment GST-3.1-3.5 in (A)). Data relates to Fig. 5.

NHSL3 contains at least 11 putative SH3 domain binding sites according to the bioinformatics tool Scansite using its most stringent settings (https://scansite4.mit.edu). Given the known SH3 domain in the Abi subunit of the Scar/WAVE complex, we tested direct interaction between NHSL3 and Abi and mapped the binding site using a far-western approach. We purified six GST-tagged NHSL3 fragments (GST1-6) from *E. coli* spanning the entire amino acid sequence (Fig. 5E; S4A,B). As a positive control, we purified a fragment of Lpd harbouring three SH3 binding sites (GST-Lpd) known to bind to Abi^24^ (Fig. 5E; S4A,B). After separation by SDS-PAGE, we blotted these proteins onto a PVDF membrane and overlayed these with either purified maltose binding protein (MBP)-tagged Abi1 (MBP-Abi), MBP-tagged Abi1 with a deletion of its SH3 domain (MBP-AbiΔSH3), MBP-only (MBP) as a negative control. This analysis revealed that Abi1 only bound to fragment 4 of NHSL3 and that this interaction required the Abi1 SH3 domain (Fig. 5D,E). To further delineate the binding site, we purified five overlapping sub-fragments (GST-3.1 to 3.5) spanning the original GST fragment 3 of NHSL3 (Fig. 5E; S4A,C). Overlay of purified MBP-Abi1 or MBP-Abi1ΔSH3 showed an SH3 domain-specific interaction only with NHSL3 sub-fragments GST-3.2 and -3.3 (Fig. 5E,F). The overlap between these two sub-fragments harbours only one putative SH3 binding site which is composed of three overlapping motifs, one SH3 class I and two SH3 class II sites (Fig. 5E; S4A). We mutated this putative site completely (by introducing point mutations into all three motifs) in full-length NHSL3 (NHSL3ΔSH3-EGFP) and tested for a loss of interaction with Abi in a co-immunoprecipitation. We co-transfected HEK cells with Myc-Abi1 and either wild-type NHSL3-EGFP (as positive control), NHSL3ΔSH3-EGFP, or NHSL3ΔFP4-1/2-EGFP. Myc-Abi1 was pulled down equally strongly by both wild-type NHSL3 (NHSL3-EGFP) and the Ena/VASP binding mutant (NHSL3ΔFP4-1/2-EGFP). This suggests that the interaction is not mediated through an indirect interaction between Abi1 and Ena/VASP proteins. The inability of the mutant NHSL3ΔSH3-EGFP to pulldown Myc-Abi indicates that the SH3 binding site in fragment 3 is necessary for the interaction with Abi1 (Fig. 6A).

**Figure 6.**
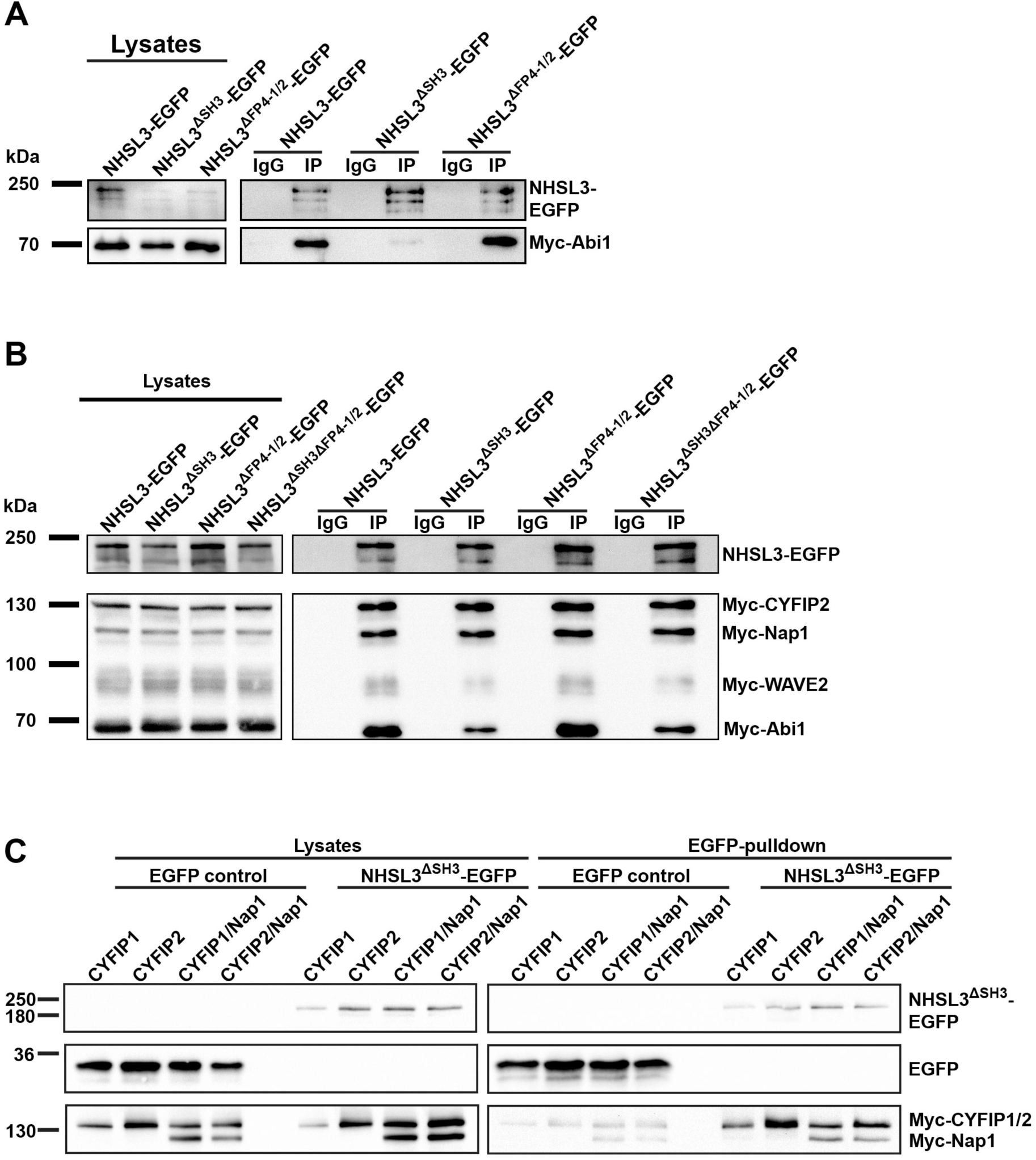
NHSL3 interacts, in addition to Abi, with the CYFIP1/2 subunits of the Scar/WAVE complex. (**A**) Myc-tagged Abi1 was co-expressed with wild-type NHSL3-EGFP, EGFP-tagged NHSL3 with the Abi SH3 domain binding site mutated (NHSL3^ΔSH3^-EGFP) or the first and second Ena/VASP binding site mutated (NHSL3^ΔFP41/2^-EGFP). EGFP-tagged NHSL3 was immunoprecipitated from cell lysates using NHSL3 polyclonal antiserum or normal rabbit IgG as negative control and blots were probed using antibodies against Myc and EGFP. Blot representative of three independent experiments. (**B**) Myc-tagged CYFIP2, Nap1, Scar/WAVE2, Abi1, and HSPC300 were co-expressed with wild-type NHSL3-EGFP, EGFP-tagged NHSL3 with the Abi SH3 domain binding site mutated (NHSL3^ΔSH3^-EGFP) or the first and second Ena/VASP binding site mutated (NHSL3^ΔFP41/2^-EGFP) or both NHSL3^ΔSH3ΔFP41/2^-EGFP. EGFP-tagged NHSL3 was immunoprecipitated from cell lysates using NHSL3 polyclonal antiserum or normal rabbit IgG as negative control and blots were probed using antibodies against Myc and EGFP. Blot representative of three independent experiments. **(C)** HEK cells were co-transfected with Myc-tagged CYFIP1, or CYFIP2, or CYFIP1 with Nap1 or CYFIP2 with Nap1 and with EGFP only as control or EGFP-tagged NHSL3 with the Abi SH3 domain binding site mutated (NHSL3^ΔSH3^-EGFP). EGFP-tagged NHSL3 was pulled down from cell lysates using a nanobody against EGFP and blots were probed using antibodies against Myc and EGFP. Representative blots from three independent experiments (CYFIP1), two independent experiments (CYFIP2), and 6 independent experiments (CYFIP1/NAP1 and CYFIP2/NAP1).

### NHSL3 binds to the CYFIP1/2-Nap1 dimer in addition to Abi1

We co-transfected HEK cells with either wild-type NHSL3-EGFP or NHSL3 mutated in the Abi or Ena/VASP binding sites (NHSL3ΔSH3-EGFP or NHSL3ΔFP4-1/2-EGFP) or both (NHSL3ΔSH3ΔFP4-1/2-EGFP) to verify that the interaction with the entire Scar/WAVE complex was lost in the Abi binding mutant, and to account for any potential indirect interaction with the Scar/WAVE complex via the known interaction between Abi and the Ena/VASP proteins^49^. We observed a reduction of interaction with Abi in the NHSL3 constructs containing mutations in the Abi binding site. Surprisingly however, we did not observe a loss of interaction as a result of any of these mutations (Fig. 6B), suggesting that the interaction between NHSL3 and the Scar/WAVE complex is not solely mediated by the Abi SH3 domain interaction. This experiment also showed that the interaction is not facilitated by an indirect interaction via Ena/VASP proteins and Abi.

The Scar/WAVE complex is composed of a dimer of CYFIP1/2-Nap1 and a trimer of Abi-Scar/WAVE-HSPC300^15^. As the remaining interaction with the CYFIP1/2-Nap1 dimer appeared strongest (Fig. 6B), we explored an alternative interaction with these. We co-transfected NHSL3ΔSH3-EGFP (to prevent interaction via Abi) with Myc-tagged CYFIP1, CYFIP2, or CYFIP1+Nap1, or CYFIP2+Nap1 into HEK cells and pulled down NHSL3-EGFP from lysates. This revealed that NHSL3ΔSH3-EGFP can interact with CYFIP1 or 2 in the CYFIP1/2-Nap1 dimer (Fig. 6C). However, from these experiments we cannot conclude that it solely interacts with CYFIP1/2 as endogenous Nap1 may form a dimer with Myc-tagged CYFIP1/2. Transfection and pulldown experiments with only Myc-Nap1 were inconclusive as Myc-Nap1 expressed on its own was sticky to the EGFP-nanobody beads (not shown), potentially due to the access of interfaces normally involved in dimer formation with CYFIP1/2.

To map this interaction in more detail, we created four overlapping EGFP-tagged fragments of NHSL3 (Fig. 7A), to minimise the risk of destroying potential unknown binding sites, and co-transfected them with Myc-tagged CYFIP1 and Nap1. Full length EGFP-tagged NHSL3 with a mutated SH3 domain binding site (NHSL3ΔSH3-EGFP) served as the positive control. We observed that fragment 1 and 2 strongly, and fragment 4 weakly pulled-down Myc-tagged CYFIP1 and Nap1 (Fig. 7A,B; quantification in Fig. 7C,D). Similar results were obtained for the CYFIP2-Nap1 dimer (Fig. S5A,B; quantification in Fig. S5C,D). This suggests that there are at least two binding sites within NHSL3 for CYFIP1/2-Nap1 and that these are within the first 486 amino acids of NHSL3. This conclusion follows from fragments 1 and 2 independently pulling-down CYFIP1/2-NAP1 but not fragment 3 which therefore excludes the latter half of NHSL3 from containing the binding site (Fig. 7A).

**Figure 7.**
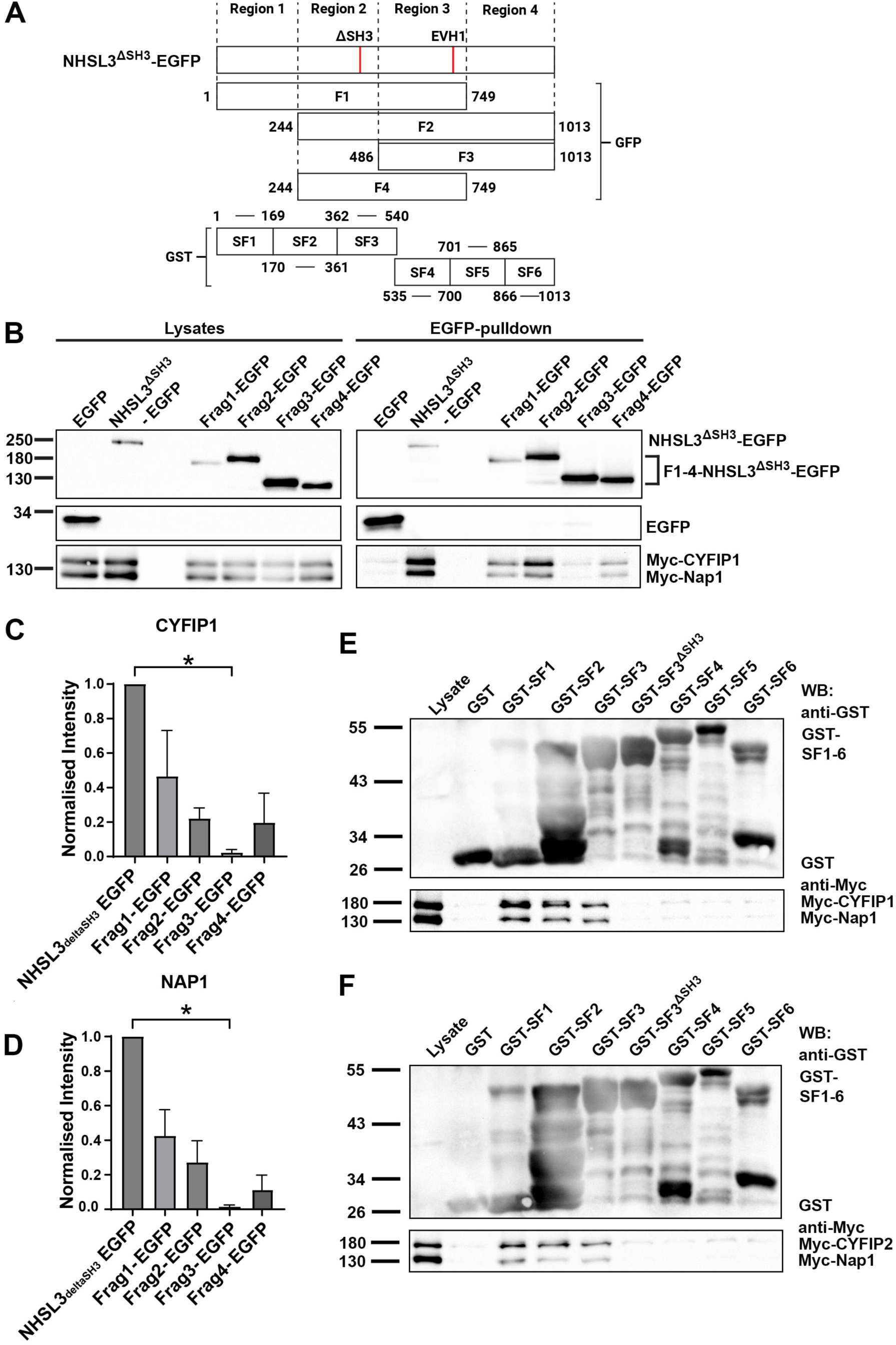
The N-terminus of NHSL3 interacts with the CYFIP1-Nap1 subunits of the Scar/WAVE complex. **(A)** Schematic overview of the location of EGFP tagged overlapping NHSL3 fragments 1-4 and the GST-tagged NHSL3 fragments 1-6 in full length NHSL3^ΔSH3^-EGFP. The location of the mutated Abi binding site and the two Ena/VASP EVH1 domain binding sites (EVH1) are depicted. The numbering refers to the amino acid numbers at the start and end of each fragment referring to murine NHSL3 isoform c (mmNHSL3 transcript variant 3 with exon1c; Genbank No: NM_001285866). **(B)** HEK cells were co-transfected with Myc-tagged CYFIP1 and Nap1 and with EGFP only as control or EGFP-tagged NHSL3 with the Abi SH3 domain binding site mutated (NHSL3^ΔSH3^-EGFP) or with NHSL3-fragment 1-4 (for location see (A)). EGFP-tagged NHSL3 proteins were pulled down from cell lysates using a nanobody against EGFP and blots were probed using antibodies against Myc and EGFP. Representative blots from three independent experiments. **(C,D)** Quantification of the EGFP-trap pulldown of CYFIP1 (C) and NAP1 (D) with NHSL3-EGFP fragments from (B). All measurements were normalised to the expression of CYFIP 1 or NAP1 and the amount of pulled down EGFP proteins. The results are displayed relative to the intensity of the pulldown of full-length NHSL3^ΔSH3^-EGFP. Data taken from three independent experiments and plotted as a bar graph, mean ± SEM. Kruskal-Wallis with Dunn’s multiple comparisons test: ∗P < 0.05. **(E,F)** HEK cells were co-transfected with Myc-tagged CYFIP1 and Nap1. Proteins from cell lysates were pulled down using purified, bead-immobilised GST-tagged NHSL3 fragments 1-6 (for localisation in full length NHSL3 see A) and blots were probed using antibodies against Myc and GST. Representative blots from three independent experiments.

**Figure S5.**
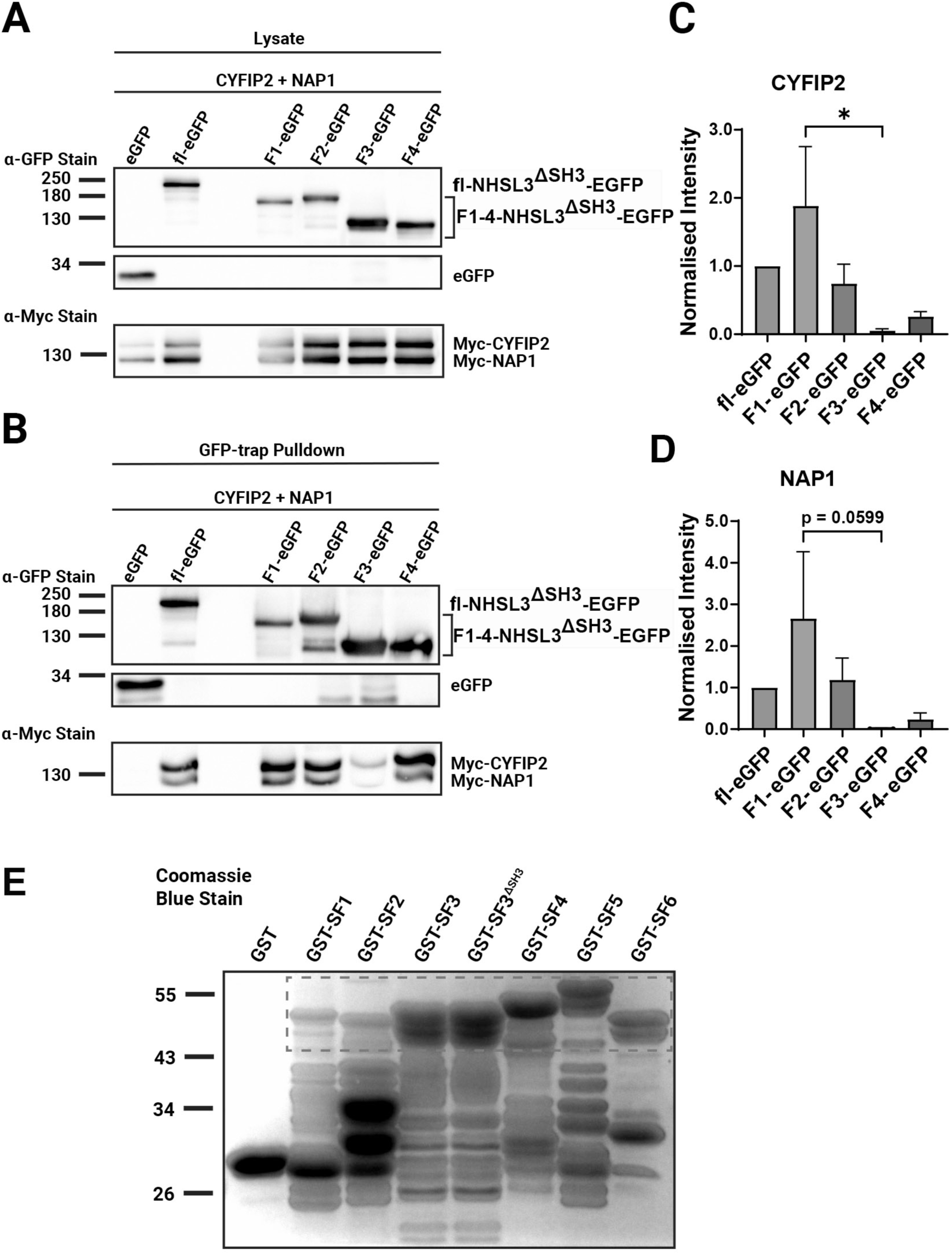
The N-terminus of NHSL3 interacts with the CYFIP2-Nap1 subunits of the Scar/WAVE complex. **(A,B)** HEK cells were co-transfected with Myc-tagged CYFIP2 and Nap1 and with EGFP only as control or EGFP-tagged NHSL3 with the Abi SH3 domain binding site mutated (NHSL3^ΔSH3^-EGFP) or with NHSL3-fragment 1-4 (for location see (Fig.7 A)). EGFP-tagged NHSL3 proteins were pulled down from cell lysates using a nanobody against EGFP and (A) lysates blots and (B) pulldown blots were probed using antibodies against Myc and EGFP. Representative blots from three independent experiments. **(C,D)** Quantification of the EGFP-trap pulldown of CYFIP2 (C) and NAP1 (D) with NHSL3-EGFP fragments from (B). All measurements were normalised to the expression of CYFIP2 or NAP1 and the amount of pulled down EGFP proteins. The results are displayed relative to the intensity of the pulldown of full-length NHSL3^ΔSH3^-EGFP. Data taken from three independent experiments and plotted as a bar graph, mean ± SEM. Kruskal-Wallis with Dunn’s multiple comparisons test: *P < 0.05. **(E)** Coomassie gel of purified GST and the GST-tagged fragments 1-6 and fragment 3 mutated in the Abi SH3 domain binding site. Please note since NHSL3 is intrinsically unstructured, protein degradation is apparent. Data relates to Fig. 7.

To further delineate the binding sites, we then re-employed the six purified GST-tagged NHSL3 fragments covering the entire NHSL3 amino acid sequence. In addition, we also purified GST-tagged fragment 3 mutated in the Abi SH3 domain binding site to prevent indirect interaction via Abi (Fig. 7A, S5E). Bead immobilised, purified GST-NHSL3 fragment fusion proteins were used to pulldown Myc-tagged CYFIP1 and Nap1 (Fig. 7E) or Myc-tagged CYFIP2 and Nap1 (Fig. 7F). This experiment showed that the CYFIP1/2-Nap1 dimer binds strongly to NHSL3 fragments 1 and 2 but to fragment 3 only in the presence of the intact Abi SH3 domain binding site suggesting that the binding sites for CYFIP1/2-Nap1 reside within the NHSL3 region spanned by fragments 1 and 2.

Taken together NHSL3 binds to CYFIP1/2 through at least two binding sites located within a 361-amino acid N-terminal region of NHSL3 (Fig. 7A). One of these sites must be located within the first 169 amino acids (SF1) since this fragment is able to bind independently of any other. The other site must be located within a 117-amino acid region (aa 244-361) starting at the beginning of the larger overlapping fragment 2 (F2) and finishing at the end of SF2. This is deduced since SF2 can bind independently of any other fragment, but F2 which only overlaps with SF2 by 117 residues is also able to strongly interact with CYFIP1/2.

### Bioinformatics and structural modelling predict three CYFIP1 binding sites in NHSL3

We employed multiple sequence alignment by fast Fourier transform (MAFFT)^50^ to determine highly conserved residues across a diverse range of species: mouse, turtle, zebrafish, chicken, human, and frog. These sequences were compared to a consensus sequence constructed using a > 70% cut-off threshold. We then identified all the residues in the mouse (*Mus musculus*) NHSL3 isoform 1c (used throughout this work) that matched this consensus sequence. The result is shown as the upper sequence (Fig. S6A). To increase species coverage, we made use of deep-learning assisted multiple-sequence alignment (DeepMSA)^51^, an open-source tool utilising homologous sequences and alignments created from whole-genome and meta-genome databases. An alignment depth (Nf) of 5.21643 and 334 sequences was reported by the analysis and the result is shown as the lower sequence (Fig. S6A) with larger letters indicating greater conservation. There are several motifs with a high level of agreement between the MAFFT and the DeepMSA results which reside within the putative binding regions identified by the NHSL3 fragment pulldown assays (Fig. S6B).

NHSL3 is mostly unstructured with a few secondary structures predicted by Alphafold2 (Fig. 8A) and which we have indicated below the DeepMSA sequence (Fig. S6A). Alphafold2 was then implemented to predict secondary structures in sub-fragments 1-2 of NHSL3 (Fig. 8B). All structures predicted with confidence or high confidence were also observed in this model. Interestingly, an α-helical secondary structure not predicted in the full-length NHSL3 model is predicted in this fragment model over S117–Q132 which incorporates part of the highly conserved motif at residue 127. The highly confident region preceding the β-hairpin predicted at V306 was recaptured with equally high confidence in this fragment model.

**Figure 8.**
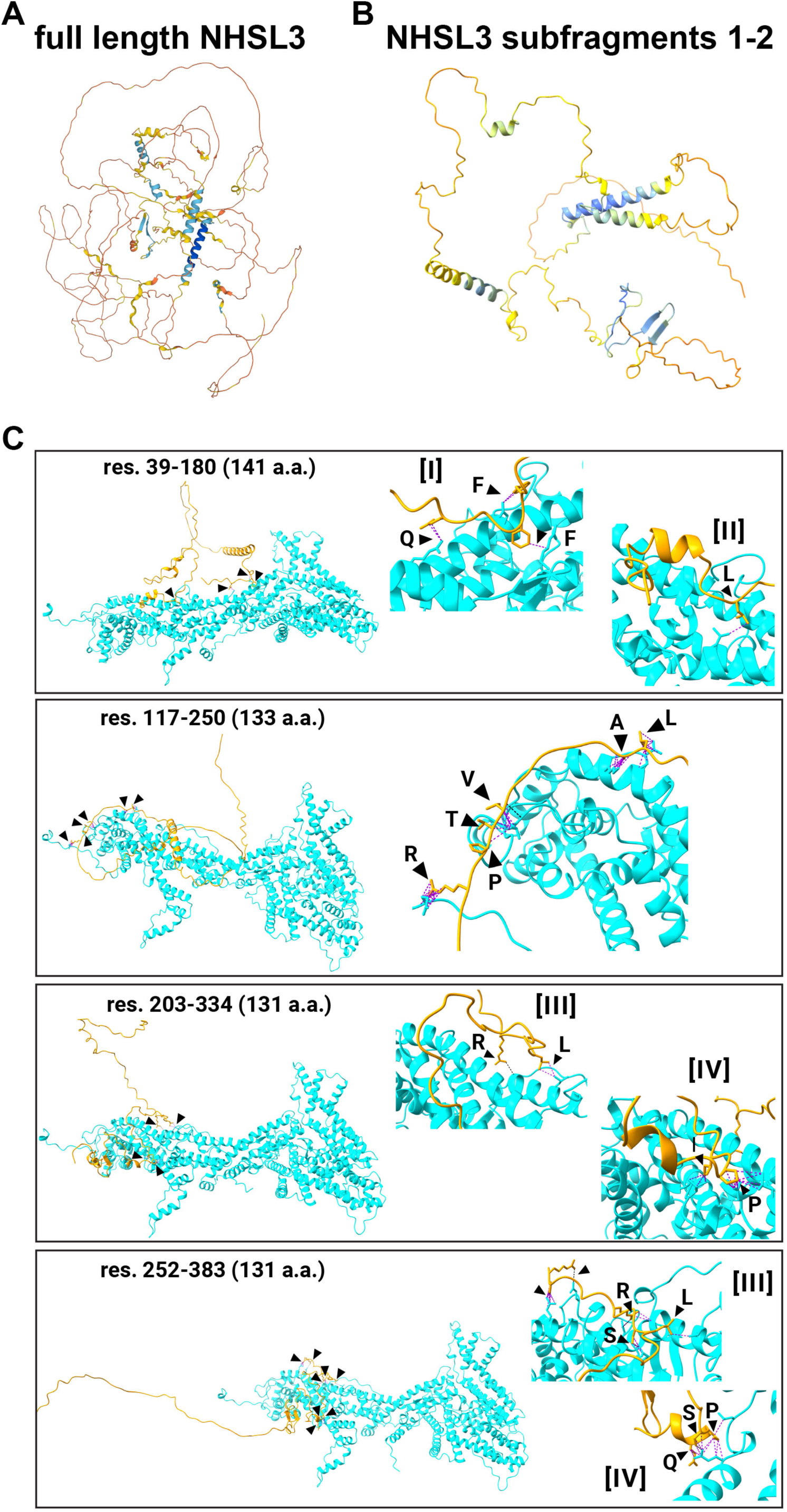
The N-terminus of NHSL3 harbours putative binding sites with the CYFIP1 subunit of the Scar/WAVE complex. (**A**) The predicted structure of full-length NHSL3 as displayed on the AlphaFold Protein Structure Database (Uniprot accession number: A2A7S8). The protein is colour coded by a per-residue confidence score (pLDDT) as on the database. (**B**) A model of the N-terminal region of NHSL3 (GST-tagged fragments SF1 and SF2 (Fig. 7A)). The structure was predicted using ColabFold and visualised in ChimeraX. The model is colour coded with the same palette as the AlphaFold database but is normalised to the range of pLDDT scores within this prediction (min = 29.8, max = 93.5). (**C**) Model predictions of four virtual overlapping fragments of the N-terminal region of NHSL3 in complex with CYFIP1 (full-length). The location and length of the fragments are given in each case (res. = residue, a.a. = amino acid). CYFIP1 is depicted in cyan, and NHSL3 fragments in orange. In each panel, the left-hand side shows an expanded view of the complex, whilst the right-hand side magnifies the regions containing points of inter-chain contact between NHSL3 and CYFIP1. In both cases arrowheads indicate the contact sites, where ball-and-stick displays of the residues in addition to the cartoon ribbon exhibit the atoms involved in the contact. Dashed magenta connecting lines indicate any van der Waals overlap, dashed black lines indicate hydrogen bonds specifically. Capital Roman numerals indicate contacts which occurred in multiple models, and which involve highly conserved sequences, labelled [I] – [IV].

**Figure S6:**
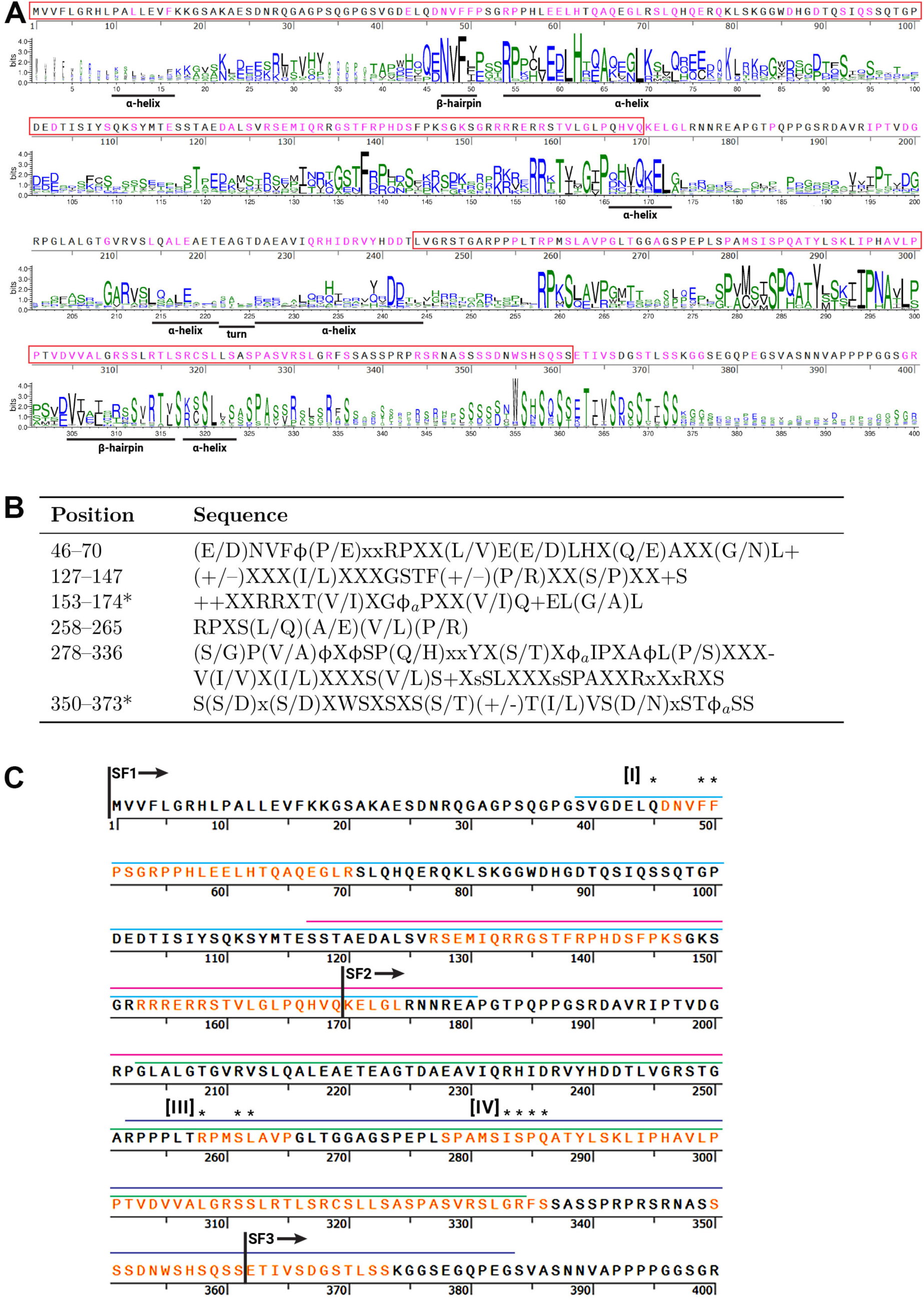
The N-terminus of NHSL3 harbours putative binding sites with the CYFIP1 subunit of the Scar/WAVE complex. **(A)** The sequence in the top row is that of mmNHSL3 isoform c, with residues conserved across vertebrate species as determined by MAFFT coloured in magenta. The binding regions identified from biochemical interaction assays (Fig. 7) are enclosed in red boxes. The second row shows the output of the DeepMSA screen with the size of the amino acid letter corresponding to the conservation at each position. The amino acids are coloured by chemical property, provided by the server (https://zhanggroup.org/DeepMSA/). Annotated underneath the DeepMSA is the secondary structure as predicted by AlphaFold2 from the protein database (Uniprot accession number: A2A7S8). **(B)** Summary of highly conserved sequences in NHSL3 based on MAFFT and DeepMSA found within the two regions identified from biochemical interaction assays (Fig. 7). X = any amino acid; x = any small amino acid; φ = a hydrophobic amino acid; φa = a hydrophobic, aliphatic amino acid; + = a positively charged amino acid; +/– = a charged amino acid. *Extended beyond the interaction regions defined by the fragment binding. **(C)** A graphical summary of conserved sequences and contact sites in the N-terminal region of NHSL3, with annotations of the real and virtual fragments used for analysis. Real fragments are indicated as SF1-3, virtual fragments are indicated by coloured lines above the sequence: cyan = 1, magenta = 2, green = 3, and dark blue = 4. Conserved sequences, as described in table (B), are indicated by orange amino acid lettering. Sites of contact are indicated by asterisks above the letter indicating the contact residue. Only contact sites that lie within conserved sequences and replicated in two fragments (where possible) are annotated. Data relates to Fig. 8.

To identify putative binding sites between the N-terminus of NHSL3 and CYFIP1, four virtual fragments of 131–141 amino acids in length were modelled in complex with CYFIP1 using Alphafold2-multimer (res. 39-180 (141 a.a.), res. 117-250 (133 a.a.), res. 203-334 (131 a.a.), res. 252-383 (131 a.a.)). The models that were ranked most highly in confidence were inspected for putative contact sites using ChimeraX and are shown in (Fig. 8C). Four motifs were identified which contained residues with contacts that were part of highly conserved sequences and were reproduced in two models where the virtual fragments overlapped. The first of these, annotated [I] in (Fig. 8C) is the 45-QDNVFFP-51 motif, in which the residues Q and FF form contacts. The second, annotated [II] in (Fig. 8C), is the 157-RRSTVLGLP-165 which may be part of a larger motif (see Table in Fig. S6B) in which just the L162 forms a contact. This is unlikely to be a real contact site, or at least not an important one, since the leucine residue is one of the least conserved in this motif, and the chemistry of this residue is not highly conserved either (the position is marked as an X in the Table in Fig. S6B). The third motif annotated [III] in (Fig. 8C) is the 258-RPMSLAVP-265 in which the R258 and L262 residues are seen to form contacts in the first model of a virtual fragment containing this sequence (third panel in the figure) and the S261 residue makes additional contact in a second model containing this sequence (fourth panel in (Fig. 8C)). The contacts further upstream of this motif seen only in the second model (fourth panel) are unlikely to be real considering that these residues are not highly conserved, and the contacts not reproduced in the first model (third panel). The final predicted binding site is 283-ISPQ-286, annotated [IV], which again, may be part of a longer motif (see Table in Fig. S6B). In the first model of a virtual fragment containing this sequence (third panel in the figure), the I283 and P285 residues form contacts. In the second model (fourth panel), S284, P285, and Q286 are in contact with CYFIP1.

Both the RPMSL site and the ISPQ site are identified in two overlapping virtual fragments giving confidence to their role in the interaction, although the exact bond-forming residues varies slightly in the two models. Only one virtual fragment contained the QDNVFFP motif, since it is so close to the N-terminus of NHSL3, and thus, we did not have a second model to corroborate this result. However, this motif is the only one present in the GST-tagged sub-fragment (SF1) that was shown in the biochemistry assays to interact with CYFIP1 (Fig. 7E), the other two putative binding motifs are both contained within SF2. It is therefore likely that this is the first binding site. The second and third sites may be either RPMSL, ISPQ, or conceivably both as the two motifs are separated by just 20 amino acids. In summary, we have identified three putative sites in NHSL3 which may mediate the interaction with CYFIP1 (Fig. S6C).

### Three sites in NHSL3 mediate the interaction with CYFIP1/2

To test the requirement of these sites for interaction of NHSL3 with CYFIP1, we mutated in total 7 amino acids covering all three sites (**Q**DNV**FF**P>**A**DNV**AA**P; **R**PMS**L**>**A**PMS**A**; **I**S**P**Q>**A**S**A**Q) in full length NHSL3 already mutated in the SH3 domain binding site (NHSL3ΔSH3-EGFP), creating the double binding-mutant NHSL3ΔSH3ΔCYFIP-EGFP. We co-transfected HEK cells with this double binding-mutant or NHSL3ΔSH3-EGFP as control together with Myc-CYFIP1 or Myc-CYFIP2. Pulldown of EGFP revealed a highly reduced interaction between NHSL3 and CYFIP1 or CYFIP2 suggesting that at least some of the predicted interaction sites mediate the binding of NHSL3 with CYFIP1/2 (Fig. 9A).

**Figure 9.**
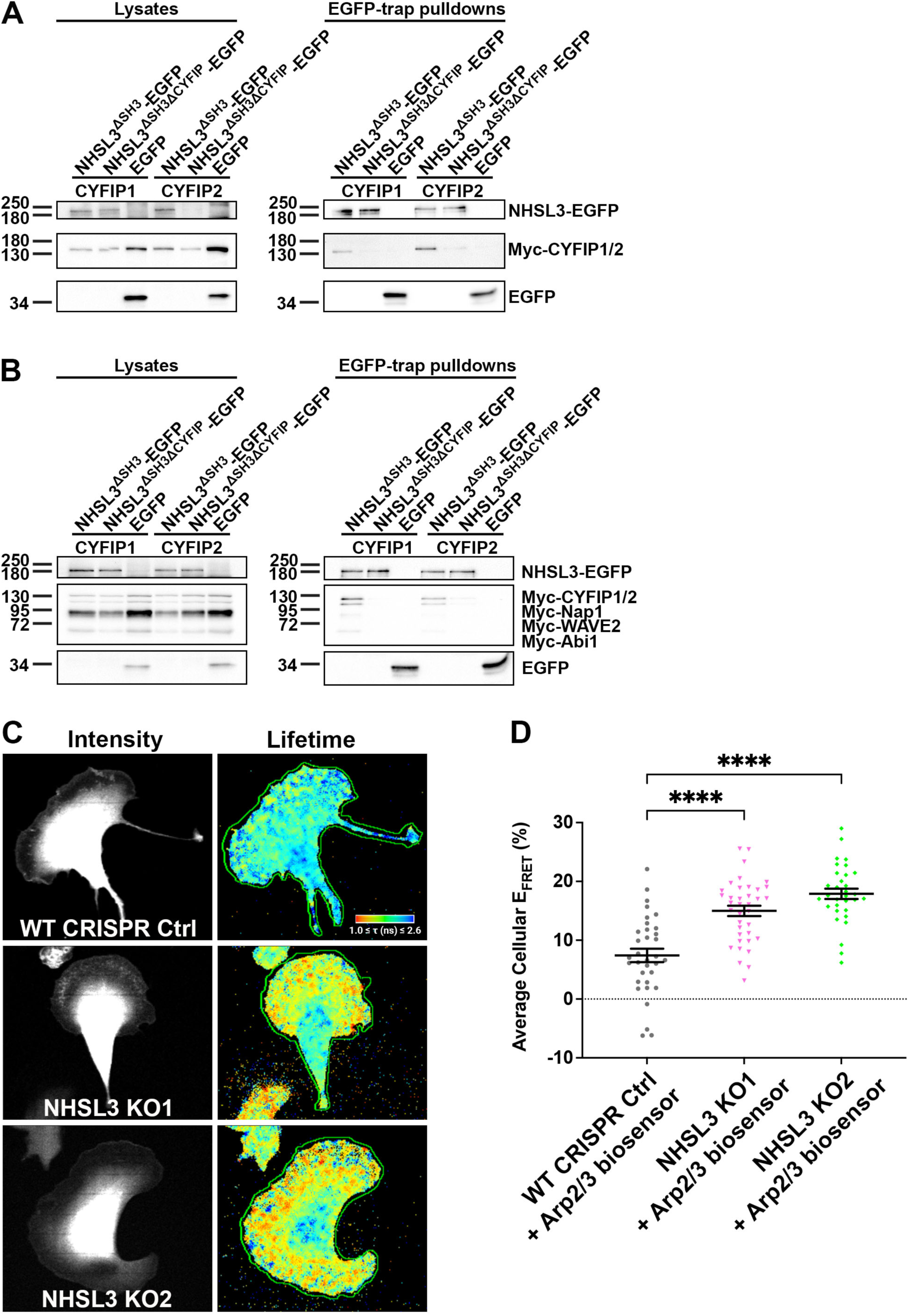
The N-terminus of NHSL3 binds to the CYFIP1 subunit via three motifs and NHSL3 is a negative regulator of the Arp2/3 complex. **(A,B)** HEK cells were co-transfected with Myc-tagged CYFIP1 or CYFIP2 (A) or Myc-tagged Nap1, CYFIP1 or CYFIP2, Nap1, Scar/WAVE2, Abi1, and HSPC300 (B) and with EGFP only as control or EGFP-tagged NHSL3 with the Abi SH3 domain binding site mutated (NHSL3^ΔSH3^-EGFP) or EGFP-tagged NHSL3 with the Abi SH3 domain binding site and the seven amino acids mediating CYFIP binding mutated (NHSL3^ΔSH3ΔCYFIP^-EGFP). EGFP-tagged NHSL3 proteins were pulled down from cell lysates using a nanobody against EGFP and blots were probed using antibodies against Myc and EGFP. Representative blots from three independent experiments. **(C,D)** Knockout of NHSL3 causes an increase in Arp2/3 complex activity. (**C**) Lifetime images of wild-type (WT) CRISPR Ctrl, NHSL3 CRISPR KO-1, and NHSL3 CRISPR KO-2 B16-F1 cells expressing the Arp2/3 complex biosensor plated on laminin coated glass. Warm colours indicate short lifetimes, and cool colours long lifetimes, of the donor fluorophore mTurq2. Shorter lifetimes represent higher FRET efficiency and therefore higher Arp2/3 complex activity. Representative images from four independent experiments. (**D**) Quantification of average cellular FRET efficiency (E_FRET_) which corresponds to Arp2/3 complex activity in WT CRISPR Ctrl (grey circles), NHSL3 CRISPR KO-1 (pink inverted triangles), and NHSL3 CRISPR KO-2 (green diamonds) cells. Data points are the weighted mean FRET efficiency across each individual cell, and bars represent the population mean ± SEM. Data from four independent experiments (N = 4): n = 34 (WT CRISPR Ctrl), n = 37 (NHSL3 CRISPR KO-1), n = 32 (NHSL3 CRISPR KO-2). Ordinary one-way ANOVA with Dunnett’s multiple comparisons test: ∗∗∗∗P < 0.0001 (WT CRISPR Ctrl <> NHSL3 CRISPR KO-1; WT CRISPR Ctrl <> NHSL3 CRISPR KO-2).

**Figure S7:**
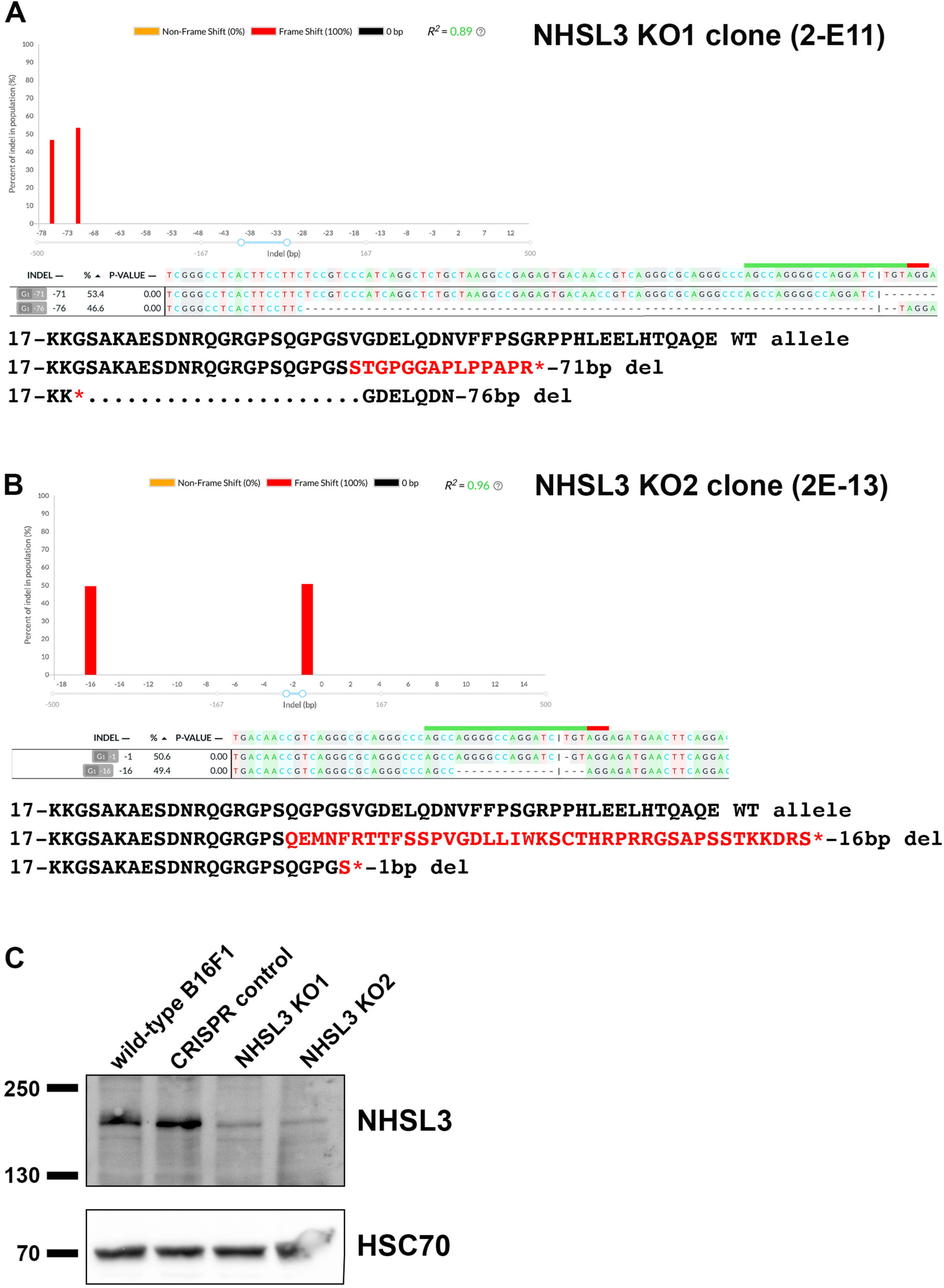
Characterisation of the NHSL3 CRISPR knockout cell lines. **(A,B)** Genomic DNA was isolated from wild-type B16-F1 cells as well as potential NHSL3 knockout clones, and the region of DNA encompassing the expected cut site in exon 2 was amplified by PCR. These amplicons were sequenced, and DNA Sequences of knock-out clones were compared to WT sequence using the DECODR web tool. **(A)** 53.4% of alleles in NHSL3 KO1 contain a deletion of 71bp or 46.6% of alleles contain a deletion of 76bp at the cut site resulting in frame shifts and premature termination codons. **(B)** 50.6% of alleles in NHSL3 KO2 contain a deletion of 1bp or 49.4% of alleles contain a deletion of 16bp at the cut site resulting in frame shifts and premature termination codons. Altered amino acid sequence as a result of the frame shift is shown in red and the premature termination codon is represented with a ‘*’. **(C)** Wild-type B16-F1 cell or the B16-F1 CRISPR control cell line or NHSL3 CRISPR knockout cell lines NHSL3 KO1 or NHSL3 KO2 were lysed and equal amounts of lysates separated by SDS-PAGE and blotted. Blots were incubated with NHSL3 (upper blot) or HSC70 antibodies as loading control (lower blot). Representative blots from three independent experiments. Data relates to Fig. 9 and 10.

To explore whether we have identified all sites required for the interaction between NHSL3 and all the components of the Scar/WAVE complex, we repeated the experiment and co-transfected HEK cells with NHSL3ΔSH3ΔCYFIP-EGFP or NHSL3ΔSH3-EGFP as a control together with all Myc-tagged components of the Scar/WAVE complex: Myc-CYFIP1 or Myc-CYFIP2, Myc-Nap1, -Scar/WAVE2, -Abi1, -HSPC300. After lysis, we pulled down EGFP-NHSL3 mutants and observed that in the NHSL3ΔSH3ΔCYFIP-EGFP mutant interaction with the CYFIP1 containing Scar/WAVE complex was virtually absent whereas the interaction with the CYFIP2 containing Scar/WAVE complex was highly reduced (Fig. 9B). Taken together, the interaction between NHSL3 and the Scar/WAVE complex is mediated by an Abi SH3 domain binding motif and several CYFIP1/2 interaction sites.

### NHSL3 is a negative regulator of the Arp2/3 complex

Since NHSL3 binds to the Scar/WAVE complex and the latter is the key activator of the Arp2/3 complex which in turn is essential for branched F-actin nucleation, we explored the effect of NHSL3 on Arp2/3 activation. For these experiments we created a NHSL3 knockout B16-F1 melanoma cell line by CRISPR-Cas9 mediated induction of indels creating premature stop codons in exon 2 common to all NHSL3 isoforms (Fig. 1D). We created clonal knockout cell lines using sgRNAs specific to NHSL3 and by testing for the absence of expression by western blot; a control cell line was also made using a Cas9 plasmid without any sgRNA. Sequencing the genomic region of exon 2 revealed that in the two clonal lines selected (NHSL3 KO1 and KO2), all alleles contained indels causing frame shift mutations (Fig. S7A,B). Western blot analysis of these NHSL3 KO1 and KO2 cell lines revealed virtual absence of NHSL3 expression. Several weak remaining bands may represent low unspecific interaction of the antiserum with other proteins which are already evident in the wild-type and control cell line (Fig. S7C).

To test the requirement of NHSL3 for Arp2/3 activation in cells, we employed our Arp2/3 biosensor validated previously^39^. This biosensor relies on the conformational change which the Arp2/3 complex undergoes upon activation^52–56^. We transfected the B16-F1 CRISPR control, NHSL3 KO1 and NHSL3 KO2 cell lines with the Arp2/3 biosensor and measured FRET by multiphoton fluorescence lifetime imaging (MP-FLIM) (Fig. 9C). Quantification of the average cellular FRET efficiency from the lifetime values, which correspond to the Arp2/3 activity, revealed that it was significantly increased in the NHSL3 knockout cell line compared to the CRISPR control cell line: (7.4 ± 1.1)% in the control to (15.0 ± 0.9)% and (17.9 ± 0.9)% in the NHSL3 KO1 and KO2 lines respectively (Fig. 9C,D). This indicates that NHSL3 is a negative regulator of the Arp2/3 complex.

### NHSL3 promotes cell migration

Given the interactions of NHSL3 with Ena/VASP proteins and the Scar/WAVE complex, key regulators of cell migration, as well as its effect on Arp2/3 complex activity, we investigated the function of NHSL3 in cell migration. Quantification of random 2D cell migration revealed that the NHSL3 KO1 and KO2 cell lines did not change their persistence but had a significantly reduced speed compared to the CRISPR control cell line (Fig. 10A,B). Overexpression of NHSL3-EGFP (Fig. S8A) displayed the expected opposite phenotype: increased speed compared to control overexpression of EGFP only and still no change in persistence (Fig. 10C,D). Taken together, this suggest that NHSL3 functions to promote cell migration.

**Figure 10.**
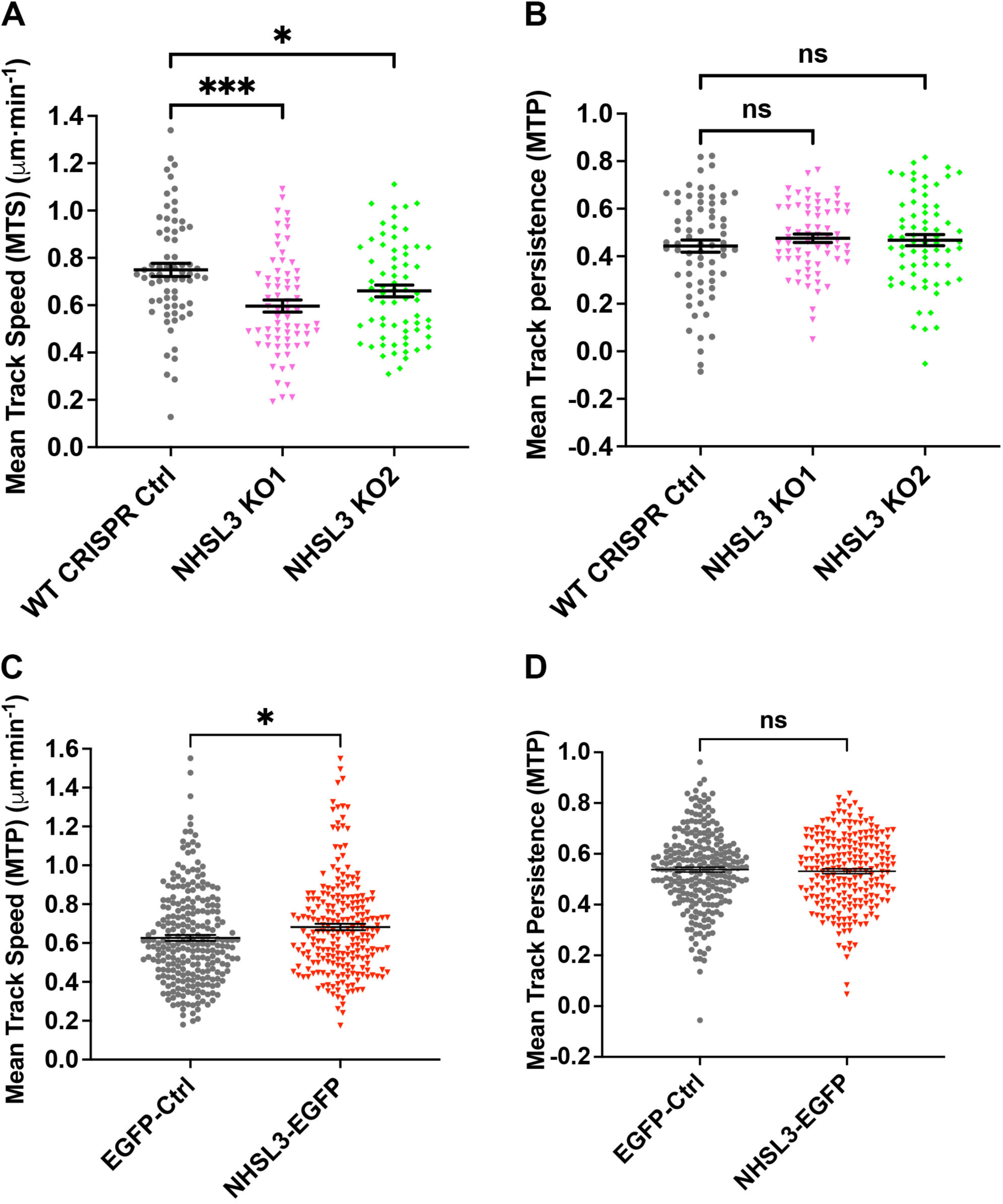
NHSL3 promotes cell migration speed but not persistence. **(A)** The mean track speed (MTS, dt = 10 mins) of randomly migrating WT CRISPR Ctrl (grey circles), NHSL3 CRISPR KO1 (pink inverted triangles), and NHSL3 CRISPR KO2 (green diamonds) B16-F1 cells plated on fibronectin. Each data point represents the MTS of an individual cell, bars represent the population mean ± SEM. Data taken from six independent experiments (N = 6): n = 69 (WT CRISPR Ctrl), n = 68 (NHSL3 CRISPR KO1), n = 68 (NHSL3 CRISPR KO2) after excluding one data point based on the identification of outliers using the ROUT method with Q = 0.1%. Kruskal-Wallis with Dunn’s multiple comparisons test: ∗∗∗P = 0.0001 (WT CRISPR Ctrl <> NHSL3 CRISPR KO1), *P = 0.0355 (WT CRISPR Ctrl <> NHSL3 CRISPR KO2). **(B)** The mean track persistence (MTP, TR = 4) of the same cells as in (A). Data points represent the MTP of each individual cell, bars represent the population mean ± SEM. Ordinary one-way ANOVA with Dunnett’s multiple comparisons test: nsP = 0.4866 (WT CRISPR Ctrl <> NHSL3 CRISPR KO1), ns P = 0.6532 (WT CRISPR Ctrl <> NHSL3 CRISPR KO2). **(C)** The mean track speed (MTS, dt = 10 mins) of randomly migrating B16-F1 cells plated on fibronectin and transiently transfected with either EGFP only as a control (EGFP-Ctrl, grey circles) or NHSL3-EGFP (red inverted triangles) to over-express NHSL3. Each data point represents the MTS of an individual cell, bars represent the population mean ± SEM. Data taken from five independent experiments (N = 5): n = 252 (EGFP-Ctrl), n = 225 (EGFP-NHSL3) after excluding two data points based on the identification of outliers using the ROUT method with Q = 0.1%. Non-parametric rank comparison (Mann-Whitney) test: *P = 0.0122 (EGFP Ctrl <> NHSL3-EGFP). **(D)** The mean track persistence (MTP, TR = 4) of the same cells as in (C). Data points represent the MTP of each individual cell, bars represent the population mean ± SEM. Unpaired t-test: ns P = 0.6613 (EGFP Ctrl <> NHSL3-EGFP).

**Figure S8:**
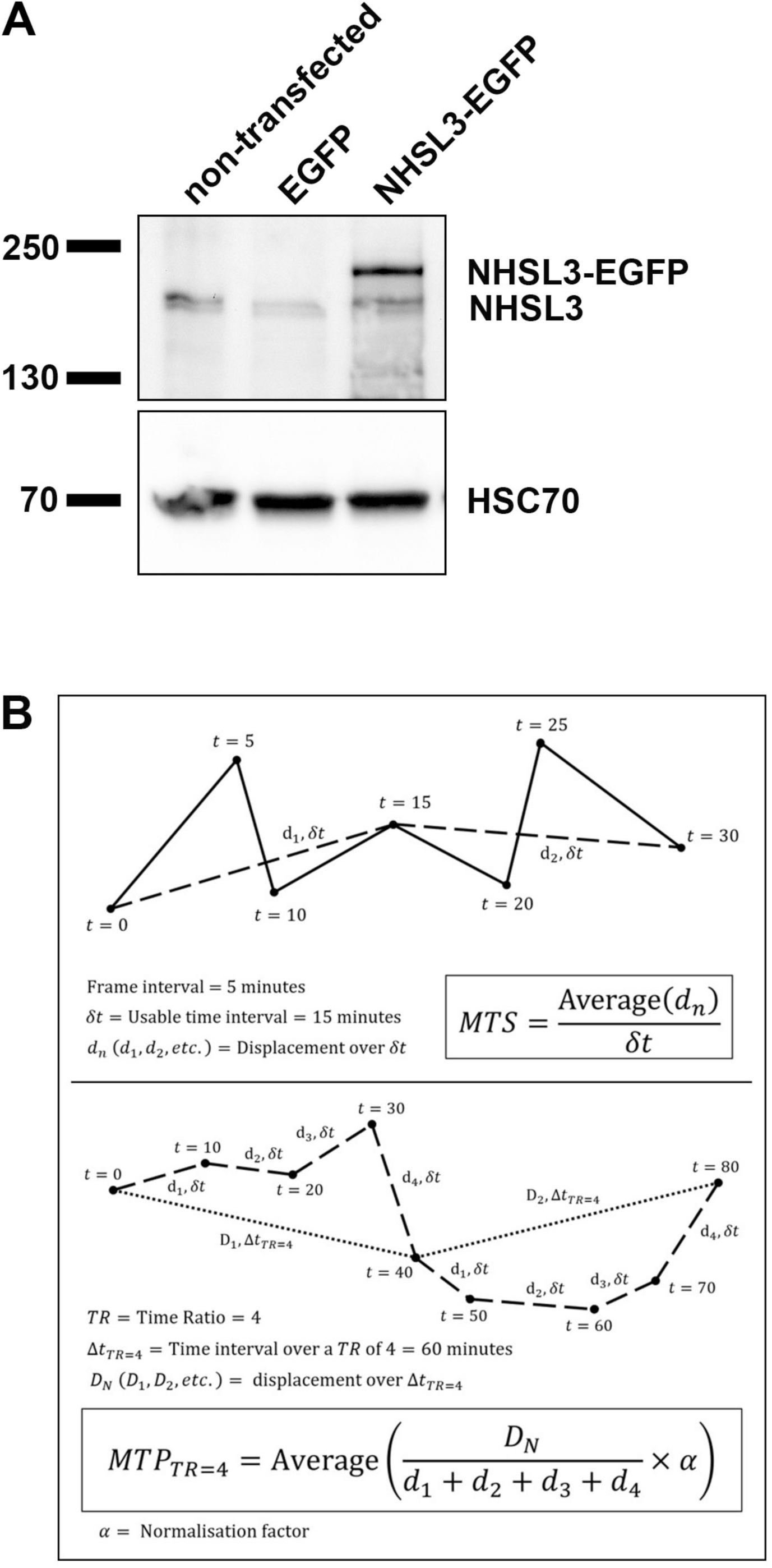
NHSL3 promotes cell migration speed but not persistence. **(A)** B16-F1 cells were either left non-transfected, transfected with EGFP only as negative control or with NHSL3-EGFP, selected by puromycin and lysed. Cell lysates were separated by SDS-PAGE and blotted. Blots were incubated with NHSL3 (upper blot) or HSC70 antibodies as loading control (lower blot). Representative blots from three independent experiments. **(B)** Schematic diagram of the definition of Mean Track Speed (MTS) and Mean Track Persistence (MTP). See detailed explanation in the methods section. Data relates to Fig. 10.

## Discussion

We identify NHSL3 as a novel protein localising to the leading edge of migrating cells. We show that NHSL3 binds to the Scar/WAVE complex which is known to control Arp2/3 mediated branched F-actin nucleation and to Ena/VASP proteins which function as F-actin elongators. These actin regulators control mesenchymal cell migration and fittingly we found that NHSL3 promotes cell migration despite acting as a negative regulator of Scar/WAVE-Arp2/3 complex activity. In contrast to other Scar/WAVE complex interactors which only bind to its Abi subunit, we revealed that NHSL3 in addition binds to its CYFIP1/2 subunit. Our findings suggest that the control of branched actin nucleation and elongation at the edge of migrating cells is even more complex than anticipated.

Our work provides evidence that KIAA1522 is indeed a novel fourth member of the NHS family, NHSL3, as interactions with the Scar/WAVE complex (observed for NHS and NHSL1) and Ena/VASP (observed at least for NHSL1^40^) are conserved. Furthermore, NHSL3 localisation to the edge of lamellipodia and vesicular structures mimics the rest of the NHS family^39–41^. Sequence similarity on the amino acid level is low, however this may be due to the overall intrinsic disorder of the proteins of the NHS family, as revealed in AlphaFold2 predictions. We tentatively suggest that only the binding motifs need to be conserved for a functional conservation, and suggest that the four members of the vertebrate NHS family may have arisen from two gene duplication events evolved from a common ancestor such as GukH in *Drosophila*^57^.

Prior to this work it has been shown that KIAA1522 (NHSL3) is upregulated in several cancers including non-small-cell lung cancer, esophageal squamous cell carcinoma, hepatocellular carcinoma, lung adenocarcinomas, colorectal carcinoma, and acute myeloid leukemia and its expression is negatively correlated with overall survival. KIAA1522 (NHSL3) has been shown to promote cellular proliferation, anoikis resistance, and consequently tumour growth and affects metastasis^45–47,58–63^. KIAA1522 (NHSL3) expression is controlled by long non-coding RNAs, circular RNAs, and miRs relevant to its role in cancer progression^63–69^. In agreement with our results that KIAA1522 (NHSL3) promotes cell migration (Fig. 10), it has been shown to increase migration in cell lines derived from hepatocellular carcinoma and colon carcinoma using scratch and transwell assays^47,58^.

Even though it is not related, Lamellipodin shows surprising similarities with proteins of the NHS family including NHSL3. Lamellipodin also binds to Ena/VASP proteins and the Scar/WAVE complex and controls cell migration^21,23–25^. Proteins of the NHS family and Lamellipodin localise to protruding lamellipodia but, in contrast to Lamellipodin, NHS family proteins do not localise to the tips of microspikes and filopodia^21,39^. The binding interfaces also differ since Lamellipodin has seven Ena/VASP domain binding sites whilst both NHSL1 and NHSL3 have only one extended site^21,40,70^. Similarly, Lamellipodin harbours three Abi SH3 domain binding sites, whilst NHSL1 has two, and NHSL3 only one binding site linking these proteins to the Scar/WAVE complex (Fig. 5,6)^24,39^. Importantly, we show here that NHSL3 differs in that it also binds to the CYFIP1/2 subunit.

Both NHSL1^39^ and NHSL3 have an overall inhibitory effect on Scar/WAVE-Arp2/3 complex activity. Despite this similarity in the regulation of branched actin, NHSL3 increases cell migration speed but has no apparent effect on persistence, in contrast to NHSL1 which functions to reduce speed and reduces persistence^39^. The effect of NHSL1 on cell migration is complex due to alternatively spliced variants. For example, both the depletion and overexpression of the NHSL1-F1 isoform increases persistence independently of any interaction with the Scar/WAVE complex^39^. But the NHSL1-A1 isoform, which can form a complex analogous to the Scar/WAVE complex (but in which Scar/WAVE is substituted for NHSL1-A1)^71,72^, resulted in a similar phenotype to NHSL3: an increase in speed without altering persistence, an effect mediated in part by an interaction with the Scar/WAVE complex^71^. It is also clear that additional functions arise in a tactic setting. We found that the NHSL1-A1 complex is required for gradient sensing during chemotaxis^71^, which supported data from Zebrafish mesodermal cells during development^73^. However, the knockdown of all NHSL1 isoforms revealed an inhibitory role in haptotaxis^72^. It is possible that these differential phenotypes are the result of the intricate balance between branching, driven by the Arp2/3 complex, and the elongation of branched filaments, driven by Ena/VASP.

We showed in pulldown assays that NHSL3, like NHSL1^40^, preferentially binds to the purified EVH1 domain of Mena, less so to VASP, and not to EVL. In contrast, Lamellipodin interacts with all Ena/VASP proteins but preferentially binds EVL^25^. However, in co-pulldown assays NHSL3 binds all three full length Ena/VASP proteins (even if it does still preferentially interact with Mena). This can be explained by tetramerisation of Ena/VASP proteins: VASP hetero-tetramerises unbiasedly with Mena, EVL, and itself. In contrast, Mena and EVL mostly homo-oligomerise^48^. An extended EVH1 domain binding site has been characterised composed of two consecutive FP4 motifs separated by approximately 15 amino acids one of which binds to the canonical binding site and the other at a second site on the opposite face of the same EVH1 domain^74^. The preference for the Mena EVH1 domain by NHSL1 and NHSL3 can be explained by such extended EVH1 domain binding sites^40^. In a genome-derived peptide interaction screen, Hwang and coworkers^31^ also identified the Mena EVH1 interaction with an extended EVH1 binding site in NHSL1, but only identified the latter half of the extended site in NHSL3. They suggested this corresponds to a weak, non-canonical motif as it only comprises DFPPPEE compared to the canonical motif F/Y/W/LPXΦP (X any and Φ hydrophobic amino acids) flanked by acidic amino acids^28–30^. Our data that mutating each site individually already causes complete loss of interaction (Fig. 4F) suggest that indeed the two motifs connected by 13 amino acids represent such an extended binding site which engages a single EVH1 domain. The first motif is canonical and is the first Ena/VASP ligand harbouring a tryptophan in the first position, which has been shown in *in vitro* binding assays to confer a higher affinity^28,29^. The second motif, 14 amino acids downstream is the formerly identified non-canonical site.

Mass spectrometry screens flagged NHSL3 as a potential interactor of the Scar/WAVE complex^75–77^ and one screen^75^ suggested from alignments that the two Abi SH3 binding sites in NHSL1 which we identified^39^ may be conserved in NHSL3. Here we showed that the first site in NHSL1 is indeed conserved in NHSL3 as an Abi SH3 domain binding site. However, the equivalent second site in NHSL3 does not interact with the SH3 domain of Abi as shown in our far western experiment since the GST-NHSL3-4 fragment does not interact (Fig. 5D). Moreover, mutating just the single identified site results in loss of interaction with Abi (Fig. 6A).

We showed (Fig. 6A,B) that the NHSL3-Scar/WAVE complex interaction is not mediated indirectly via Abi binding to Ena/VASP proteins. In addition to Abi, we found by Alphafold modelling that the NHSL3-Scar/WAVE complex interaction is mediated by a novel interaction with seven amino acids contained within three motifs in CYFIP1 and verified this by mutation. However, from our analysis we cannot be certain whether all seven amino acids are necessary for this interaction.

In the first interaction site, three residues contact three conjugate residues (Fig. 8C) beginning just over half-way through CYFIP1. The second site involves two motifs separated by twenty amino acids, which may form a more complex three-dimensional interaction interface with a 250-amino acid stretch in CYFIP1 more proximal to the C-terminus. These three motifs are surface accessible in the full Scar/WAVE complex and not occluded by Nap1 in the CYFIP1/2-Nap1 dimer. Furthermore, these motifs are not overlapping with the two Rac binding sites^19^ on CYFIP1/2. However, since these motifs are on the same side of the CYFIP-Nap1 dimer as the WCA binding sites, and the Scar/WAVE complex is auto-inhibited through an intramolecular interaction of the Scar/WAVE WCA domain with a CYFIP1/2-Nap1 interface, NHSL3 binding may cause and/or stabilise the active conformation of the Scar/WAVE complex.

In summary, we characterise in this study the fourth member of the NHS family as a novel lamellipodial protein and interactor of the key actin effectors: Ena/VASP proteins and the Scar/WAVE complex. NHSL3 controls Arp2/3 activity and promotes cell migration by increasing speed. This suggests that the controlled balance between branched actin nucleation and elongation at the edge of migrating cells is even more complex than anticipated.

## Materials and methods

### Molecular biology, plasmids, and reagents

Full length mouse NHSL3 isoform c cDNA including exon 1c was generated from EST clone BE309569 adding the missing first 10 amino acids in exon 1c of NHSL3 isoform 1c (1013 aa) by including the corresponding bases into the 5’ primer and cloning it into pENTR3C (Invitrogen) without a stop codon. Fragments of NHSL3 were cloned into pENTR3C (Invitrogen). NHSL3 cDNA in pENTR was mutated using Quikchange (Agilent) to create the Abi SH3 domain binding site mutant (aa433-MKR(P>A)(P>A)P(P>A)(P>A)RRTY-aa444) and the EVH1 domain binding site mutants (aa707-S(W>A)(P>A)(P>A)(P>A)PPPAPEEQDLSMAD(F>A)(P>A)(P>A)PEEV-aa732). The 7 amino acid mutations to disrupt the CYFIP binding sites were cloned by NEB HIFI cloning using a synthesized mutated N-terminal fragment (gBlock, IDT): Site 1: aa42-(DEL(Q>A)DNV(F>A)(F>A)PSG-aa53; Site 3: aa255-PLT(R>A)PMS(L>A)AVPGLT-aa268); Site 4: aa280-AMS(I>A)S(P>A)QAT-aa288 into pENTR-3C-NHSL3 to replace 908bp from the wild-type N-terminus which was removed by restriction digest.

The tagged Ena/VASP protein constructs were made using murine cDNA for VASP (gene bank no: X98475), Mena (gene bank no: U72520), EVL (gene bank no: U72519) kindly provided by Frank Gertler, MIT, and cloned into pENTR-3C with a stop codon. The tagged Scar/WAVE complex components and the Arp2/3 biosensor was described in^39^. In brief: ARPC1B (HsCD00370547), ARPC3 (HsCD00288065), Sra1 (CYFIP1; HsCD00042136), PIR121 (CYFIP2; HsCD00045545), Nap1 (NCKAP1; HsCD00045562), Abi2 (HsCD00042752), and HSPC300 (C3orf10; HsCD00045008) cloned into pENTR233 or pDONR221 (Harvard Institute of Proteomics). pDONR221-WAVE-2 (German Resource Centre for Genome Research). hsAbi1d (BC024254; Geneservice) full-length and Abi1d-ΔSH3 (aa 1–417) were cloned into pENTR11. ARPC1B-mVenus-P2A-ARPC3-mTurq2 was generated by Gibson assembly (NEB Hifi, New England Biolabs, Inc.) in pENTR3C (Invitrogen). pENTR3C-ARPC1B-mVenus-P2A-ARPC3-mTurq2 was transferred to pRK5-Myc-DEST for CMV driven expression in mammalian cells.

cDNAs in pENTR or pDONR were transferred into tagged mammalian expression vectors using Gateway^®^ recombination (Invitrogen): pCAG-DEST-EGFP, pCAG-DEST-EGFP-T2A-Puro, pCAG-EGFP-DEST-IRES-Puro, pRK5-Myc-DEST, pCAG-Myc-DEST-IRES-Puro, pCAG-Myc-DEST-IRES-BLAST, pCAG-Myc-DEST-IRES-Puro-T2A-LifeAct-EGFP, pCDNA3.1-mScarlet-I-DEST, pV16-gateway (MBP) which were generated by traditional restriction-based cloning or Gibson assembly (NEB Hifi, New England Biolabs, Inc.) cloning.

The six fragments covering the entire length of NHSL3 and the five sub-fragments of NHSL3-GST3 (see Fig. 5E for boundaries) and the EVH1 domains of VASP (1-114), Mena (1-112), and EVL (1-113) were cloned by PCR and restriction cloning into pGEX-6P1 (Amersham) or pENTR3C and transferred using Gateway^®^ recombination into pDEST15 (Invitrogen) and purified from *E.coli* on Glutathione-sepharose (Amersham). All constructs were verified by sequencing.

### Antibodies

Immobilised NHSL3 GST-4 (aa535-701; see Fig. 5E) (pAb) was digested with Prescission protease (Amersham Pharmacia Biotech) and used to raise polyclonal rabbit antiserum #3915 (Covance). Commercial primary antibodies: EGFP (Roche), Myc (9E10, Sigma), MBP (New England Biolabs), Abi1 (MBL, clone 1B9, D147-3), Scar/WAVE1 (BD 612276), Scar/WAVE2 rabbit mAb (D2C8, CST 3659), Mena mAb clone A351F7D9 (Lebrand et al., 2004); HSC70 (Santa Cruz sc7298). Secondary antibodies: HRP-anti-rabbit, -anti-mouse (Dako or CST).

### Northern blot

The murine multiple tissue northern blot (Ambion) was probed with a 680 bp NHSL3 fragment covering exon 5 and the beginning of exon 6 common to all isoforms (the last 34 amino acids of fragment GST-1 and all of GST-2) (Fig. 5E) according to manufacturer’s instructions.

### Generation of CRISPR cell lines

Stable CRISPR NHSL3 knockout B16-F1 cell lines were made by creating indels into exon 2, which is common to all isoforms. We utilised CRISPR-Cas9: Two NHSL3 specific sgRNA (sgRNA1: caccgGCCCTGACGGTTGTCACTCT; sgRNA2: caccgAGCCAGGGGCCAGGATCTGT) were designed using the Zheng lab CRISPR sgRNA design tool https://www.zlab.bio/resources and cloned into pX330A1×2, harboring Cas9. pX330A-1×2^78^ was a gift from Takashi Yamamoto (Addgene plasmid # 58766; http://n2t.net/addgene:58766; RRID:Addgene_58766). B16-F1 cells were transiently transfected with Cas9 plasmid including sgRNA’s or empty plasmid to create a control cell line and clonal cell lines were generated by limited dilution and loss of expression of NHSL3 tested in a western blot. Two NHSL3 knockout cell lines, 2E-11 and 2E-13 and the control cell line 1269-14 were selected. Genomic DNA was extracted using Qiagen QIAamp DNA mini kit and knockout cell lines genotyped using primers amplifying exon 2: (primer 1: GTGTGGTCCTATTATGTGATG; primer 2: CCAGAGGCTAAGTCAGAAGG) and sequenced by the sanger method on both strands. Indel formation was analysed using DECODR (https://decodr.org/) analysis.

### Cell culture and transient transfections

HEK293FT cells (Thermo Fisher Scientific R70007) and B16-F1 mouse melanoma cells (ATCC CRL-6323) were cultured in Dulbeccos modified Eagle’s medium containing penicillin/streptomycin, L-glutamine and 10% fetal bovine serum. Transient transfections in HEK293FT cells were carried out using Lipofectamine 2000 (Invitrogen) according to manufacturer’s instructions or PEI (Sigma 913375 or 408727: add 2ug DNA in 100ul OptiMEM to 4ul 2mg/ml PEI in 100ul OptiMEM, mix and incubate for 20 min room temperature). Transient transfections with B16-F1 cells were carried out with X-tremeGene 9 (Roche) and replaced with normal growth media 4-6 hours after transfection. All cells were maintained at 37°C in 10% CO_2_.

### Immunoprecipitations, Pulldowns and Western Blotting

Cells were lysed in glutathione S-transferase (GST) buffer (50mM Tris-HCL, pH 7.4, 200 mM NaCl, 1% NP-40, 2 mM MgCl_2_, 10% glycerol, NaF +Na_3_VO_4_, complete mini tablets without EDTA, Roche). Lysates were incubated on ice for 15 minutes and centrifuged at 17,000x*g* at 4°C for 10 minutes. Protein concentration was then determined (Pierce BCA protein assay kit; Thermo Fisher Scientific). For pulldowns, the lysate was incubated with either glutathione beads (Amersham), GFP-trap or Myc-trap beads (Chromotek) or GFP-selector beads (NanoTag). For immunoprecipitations, Protein A bead precleared lysates were incubated with primary antibody or non-immune control IgG followed by 1% BSA blocked protein A beads (Pierce) or protein A/G beads (Alpha Diagnostics). For GFP-trap, GFP-selector, or Myc-trap pulldowns, beads were blocked with 1% BSA before incubating with lysates for 1-2 hours or 30 mins, respectively. Following bead incubation, all beads were washed with lysis buffer, separated on SDS-PAGE gels and transferred onto Immobilon-P membranes (EMD Millipore). Western blotting was performed by transferring at 100 V, 350 mA, 50 W for 1.5 hours before blocking in 5% BSA, 5% or 10% milk overnight followed by 1 hour incubation with the indicated primary antibodies followed by HRP conjugated secondary antibodies (Dako or CST) for 1 hour at room temperature. Blots were developed with the Immun-Star WesternC ECL kit (Bio-Rad Laboratories) using the Bio-Rad Imager and ImageLab software.

### Protein purifications and far-western blot

GST and MBP fusion proteins were purified from BL21-CodonPlus (DE3)-RP *E. coli* (Stratagene) using glutathione (GE Healthcare) or amylose (New England Biolabs, Inc.) beads. Purified GST-NHSL3 fragments were separated on SDS-PAGE and transferred to PVDF membranes and overlayed as described previously^29^ with purified MBP-Abi full-length or MBP-Abi-delta-SH3, and MBP was detected with MBP antibodies (New England Biolabs).

### Immunofluorescence and live cell imaging

For immunofluorescence analysis cells were plated on nitric acid washed coverslips (Hecht-Assistant), coated with 25 ug/ml laminin (L2020, Sigma) and fixed 10 minutes with 4% paraformaldehyde-PHEM (60mM PIPES, 25mM HEPES, 10mM EGTA, 2mM MgCl_2_, 0.12M sucrose). For NHSL3 pAb: cells were permeabilised 2 min with 0.1% Triton-X-100/TBS and quenched for 10 min with 1 mg/ml Sodium Borohydride or cells were permeabilised with 0.05% saponin, 2 hours at RT. For all antibodies: cells were blocked with 10% normal goat serum and 10% BSA, TBS. Secondary antibodies: goat anti-rabbit, or anti-mouse Alexa488 or 568 (Invitrogen) and mounted in Prolong Diamond (Invitrogen). Cells were imaged on an IX81 Olympus microscope (see below).

For low magnification phase contrast and high magnification imaging, cells were plated on 12-well tissue culture dishes or glass bottom dishes (Ibidi; 81218-200) coated with 10 µg/ml fibronectin (F1141, Sigma) or coated with 25 µg/ml laminin (L2020, Sigma). For immunofluorescence and live imaging an IX 81 microscope (Olympus), with a Solent Scientific incubation chamber, filter wheels (Sutter), an ASI X-Y stage, Cascade II 512B camera (Photometrics), and 4× UPlanFL, 10× UPlanFL, 60× Plan-Apochromat NA 1.45, or 100× UPlan-Apochromat S NA 1.4 objective lenses) controlled by MetaMorph software was used. Supplementary movies were prepared in Metamorph and Fiji.

### Quantification of random cell migration speed and persistence

For random migration, B16-F1 cells were plated on to fibronectin coated 12-well dishes for 3 hours before imaging for 24 hours every 5 minutes at 4x magnification. Cells were manually tracked by their nuclear position using the Manual Tracking plugin (FIJI), and the cell track coordinates imported into Mathematica for analysis using the Chemotaxis Analysis Notebook v1.6β (G. Dunn, King’s College London, UK^24^).

Speed and persistence measurements from tracking data are susceptible to positional error (due to manual tracking) as well as biological noise (from cell morphology changes or nuclear repositioning). To address the former, we previously had estimated the positional error by tracking the same cell multiple times (each time blinded to the previous track) and then calculated the time interval (as an integer multiple of the frame interval) over which this error fell to at least below 10% of the average displacement measurement. The smallest multiple for which this condition was true so as to avoid error from sampling too infrequently to measure the true path length was 10 minutes^39^. This time interval is termed the ‘usable time interval’ and denoted 𝛿𝑡 (see Fig. S8B). In order to be consistent over data sets we a adopted a usable time interval that was common to all data within a dataset. This was 𝛿𝑡 = 10 (equivalent to a multiple of 2 frame intervals or taking a displacement measurement every 2 frames) for the CRISPR knockout and overexpression experiments on fibronectin. This ‘usable time interval’ approach also helps to reduce the impact of biological noise on the data.

Cell speed is the displacement (𝑑*_n_*) over 𝛿𝑡, where 𝑛 denotes which interval of the track we are measuring (for example 𝑛 = 1 would indicate the first 10 minutes, 𝑛 = 2 the second 10 minutes etc.). The Mean Track Speed (𝑀𝑇𝑆) is then the average cell speed over the whole track length (Fig. S8B, boxed equation).

Cell persistence has traditionally been defined by the directionality ratio: the net (straight line) displacement divided by the total track length. This measurement comes with additional issues to noise, namely that it is dependent on interval size, track length, and even cell speed. The interval size is determined by the usable time interval, 𝛿𝑡 for the same reasons as previously described. To address the dependence on track length, we measure persistence over another time interval, denoted Δ𝑡, which is an integer multiple of 𝛿𝑡 (see Fig. S8B, lower panel). This multiple is called the Time Ratio (𝑇𝑅, equal to Δ𝑡⁄𝛿𝑡). This approach allows tracks of different total length to be compared equally, without skewing average results.

Persistence is defined by the Mean Track Persistence (𝑀𝑇𝑃), equal to the average directionality ratio over the intervals Δ𝑡 (Δ𝑡 = 40 mins for 𝑇𝑅 = 4), multiplied by a normalisation factor, 𝛼, which sets the maximum persistence to 1 (a perfectly straight migration path) and the minimum persistence to that of a purely random walk. The directionality ratio is the net displacement, 𝐷*_N_*, where 𝑁 is the interval number for each Δ𝑡 (i.e. 𝑁 = 1 indicates the first 40 minutes, 𝑁 = 2 indicates the second 40 minutes etc.) divided by the track length over Δ𝑡 (the sum of the displacements 𝑑*_n_*) (Fig. S8B, lower panel, boxed equation).

Finally, the issue of cell speed dependence can be addressed by selecting an appropriate combination of 𝛿𝑡 and 𝑇𝑅. Plotting 𝑀𝑇𝑆 as a function of 𝛿𝑡 reveals that speed falls off exponentially with increasing time interval, for all intervals which yield displacement values above the level of track noise. The mean persistence profile is a plot of the MTP obtained for all possible choices of 𝛿𝑡 for a given 𝑇𝑅. In the mean persistence profile, the persistence normally starts off low at small speed intervals and rapidly increases to a peak before falling off again. To avoid noise, we chose a suitable 𝛿𝑡 and 𝑇𝑅 combination allowing the MTP to be measured close to this peak. The same 𝑇𝑅 must be used across all data sets so they are directly comparable; thus, another compromise must be that the 𝑇𝑅 chosen is appropriate for all populations. After some investigation, a 𝛿𝑡 of 10 and 𝑇𝑅 of 4 was chosen, for the CRISPR knockout and overexpression experiments on fibronectin. These combinations ensured in both cases that tracks were long enough to contain multiple intervals.

### FRET-FLIM analysis of Arp2/3 activity

B16-F1 cells plated on nitric acid cleaned No 1.5 coverslips (Hecht Assistent), coated with 25 ug/ml laminin (Sigma) were fixed for 20 minutes with 4% paraformaldehyde-PHEM (60mM PIPES, 25mM HEPES, 10mM EGTA, 2mM MgCl_2_, 0.12M sucrose), permeabilised for 2 minutes with 0.1% TX-100/PBS, background fluorescence quenched with 1 mg/ml sodium borohydride in PBS and mounted in ProLong Diamond (Thermo Fisher).

Time-domain FLIM data were acquired via a time-correlated single photon counting (TCSPC) custom-built, automated, 2-photon microscope. Briefly, this consisted of a Modelocked femtosecond Ti:Sapphire laser (Coherent Vision II; Coherent (UK) Ltd, Scotland) for fluorescence excitation, a dual-axis scanner, a photomultiplier detector (HPM-100-06; Becker & Hickl GmbH, Germany) and TCSPC electronics (SPC830; Becker & Hickl GmbH, Germany). Images were acquired using a 1.3 NA 40× Plan Fluor Oil Immersion objective (Nikon Instruments Ltd, UK). Fluorescence lifetimes were determined for every pixel using a modified Levenberg-Marquardt fitting technique as described previously^79^. The FLIM images were batch analysed by running an in-house exponential fitting algorithm (TRI2 software) written in LabWindows/CVI (National Instruments, Austin, TX)^79^. The fitting parameters for each time-resolved intensity image were recorded in individual output files and used to generate a distribution of lifetime and an average fluorescence lifetime. FLIM/FRET analysis was performed to investigate Arp2/3 activity using the Arp2/3 biosensor (ARPC1B-mVenus-P2A-ARPC3-mTurq2). FRET efficiencies were calculated based on the equation *E* = 1 − *τ_DA_*/*τ_D_*, where *τ_D_* and *τ_DA_* are the measured fluorescence lifetimes of the donor in the absence and presence of the acceptor, respectively.

### Statistical analyses

Data were tested for normal distribution by D’Agostino & Pearson and Shapiro-Wilk normality tests. Statistical analysis was performed in Prism 8-10 (GraphPAD Software) using a Student’s *t*-test, One-way ANOVA or non-parametric Kruskal-Wallis test with appropriate post-hoc tests (see figure legends in each case). P values <0.05 were considered significant.

## Acknowledgments

We thank Frank Gertler, MIT for reagents. Shamsinar Jalal (King’s College London, UK) for biochemistry training. F.M. was supported by a Medical Research Council (MRC) studentship. T.P. was supported by an Engineering and Physical Sciences Research Council (EPSRC) studentship. S.J. was supported by a Malaysian Public Service Department (PSD) studentship. SP is supported by UK Research and Innovation Future Leaders Fellowship (MR/T04067X/1). This work was supported by grants from the Biotechnology and Biological Science Research Council (BBSRC), UK (BB/N000226/1; BB/R015953/1) (M.K.) and the Wellcome Trust (311428/Z/24/Z) (S.A.-B).

## Data availability statement

The datasets generated or analysed during the current study are available from the corresponding author on reasonable request.

## Availability of unique biological material generated in this study

All unique biological materials such as cell lines or plasmids generated in this study are available upon reasonable request from the corresponding author.

## Supplemental movie legends

**Video S1. NHSL3 localises to the very edge of lamellipodia**

This movie shows that C-terminally EGFP-tagged NHSL3 expressed in B16-F1 cells plated on laminin localises to the very edge of protruding lamellipodia. Representative movie shown from three independent biological repeats. Cells were imaged every 10 seconds for the indicated times in minutes and seconds by wide-field time-lapse video microscopy using an IX 81 microscope (Olympus). Scale bar: 20 μm.

**Video S2. NHSL3 co-localise with the Ena/VASP family protein, Mena.**

This movie shows NHSL3-EGFP co-expressed with mScarlet-I-tagged Mena co-localise at the leading edge of cells in B16-F1 cells plated on laminin. Representative movies shown from three independent experiments. Cells were imaged every 10 seconds for the indicated times in minutes and seconds by wide-field time-lapse video microscopy using an IX 81 microscope (Olympus). Scale bar: 20 μm.

**Video S3. NHSL3 co-localise with components of the Scar/WAVE complex: Scar/WAVE2.**

This movie shows NHSL3-EGFP co-expressed with mScarlet-I-tagged Scar/WAVE2 co-localise at the leading edge of cells in B16-F1 cells plated on laminin. Representative movies shown from three independent experiments. Cells were imaged every 10 seconds for the indicated times in minutes and seconds by wide-field time-lapse video microscopy using an IX 81 microscope (Olympus). Scale bar: 20 μm.

**Video S4. NHSL3 co-localise with components of the Scar/WAVE complex: Abi1.**

This movie shows NHSL3-EGFP co-expressed with mScarlet-I-tagged Abi1 co-localise at the leading edge of cells in B16-F1 cells plated on laminin. Representative movies shown from three independent experiments. Cells were imaged every 10 seconds for the indicated times in minutes and seconds by wide-field time-lapse video microscopy using an IX 81 microscope (Olympus). Scale bar: 20 μm.

**Video S5. NHSL3 co-localise with components of the Scar/WAVE complex: Nap1.**

This movie shows NHSL3-EGFP co-expressed with mScarlet-I-tagged Nap1 co-localise at the leading edge of cells in B16-F1 cells plated on laminin. Representative movies shown from three independent experiments. Cells were imaged every 10 seconds for the indicated times in minutes and seconds by wide-field time-lapse video microscopy using an IX 81 microscope (Olympus). Scale bar: 20 μm.

**Video S6. NHSL3 co-localise with components of the Scar/WAVE complex: Scar/WAVE1.**

This movie shows NHSL3-EGFP co-expressed with mScarlet-I-tagged Scar/WAVE1 co-localise at the leading edge of cells in B16-F1 cells plated on laminin. Representative movies shown from three independent experiments. Cells were imaged every 10 seconds for the indicated times in minutes and seconds by wide-field time-lapse video microscopy using an IX 81 microscope (Olympus). Scale bar: 20 μm.

**Video S7. NHSL3 co-localise with components of the Scar/WAVE complex: Scar/WAVE3.**

This movie shows NHSL3-EGFP co-expressed with mScarlet-I-tagged Scar/WAVE3 co-localise at the leading edge of cells in B16-F1 cells plated on laminin. Representative movies shown from three independent experiments. Cells were imaged every 10 seconds for the indicated times in minutes and seconds by wide-field time-lapse video microscopy using an IX 81 microscope (Olympus). Scale bar: 20 μm.

